# A NANOG-pERK reciprocal regulatory circuit mediates Nanog autoregulation and ERK signaling dynamics

**DOI:** 10.1101/2021.06.28.450121

**Authors:** Hanuman T Kale, Rajendra Singh Rajpurohit, Debabrata Jana, Vishnu V Vijay, Mansi Srivastava, Preeti R Mourya, Gunda Srinivas, P Chandra Shekar

## Abstract

The self-renewal and differentiation potential of Embryonic stem cells (ESCs) is maintained by the regulated expression of core pluripotency factors. The expression level of core pluripotency factor *Nanog* is tightly regulated by a negative feedback autorepression loop. However, it remains unclear how the ESCs perceive the NANOG levels and execute autorepression. Here, we show that a dose-dependent induction of *Fgfbp1* and *Fgfr2* by NANOG activates an autocrine mediated ERK signaling in high-*Nanog* cells to trigger autorepression. pERK recruits NONO to *Nanog* locus to repress transcription by preventing POL2 loading. The *Nanog* autorepression process establishes a self-perpetuating NANOG-pERK reciprocal regulatory circuit. We further demonstrate that the reciprocal regulatory circuit induces the pERK heterogeneity and ERK signaling dynamics in pluripotent stem cells.

---

Embryonic stem (ES) cells are characterized by long-term self-renewal and the potential to differentiate to all cell types of the germ layers. ES cells cultured in presence of serum and LIF manifest transcriptional and functional heterogeneity. The heterogeneous expression of transcription factors like *Nanog*, *Rex1*, *Stella*, *Esrrb*, *Klf4*, and *Tbx3* determine differential fate choice(1–9). The core pluripotency factor, *Nanog* was identified as a factor conferring LIF independent self-renewal to ES cells by inhibiting differentiation(10, 11). *Nanog* switches between mono-allelic and bi-allelic expression during embryonic development and in alternate pluripotency states(3, 12). The expression of *Nanog* is restricted in ES cells to ensure their potential to differentiate by negative feedback autorepression and other repressive mechanisms(13–19). Among the multiple mechanisms that regulate *Nanog*, which mechanisms are utilized by the pluripotent cells to restrict *Nanog* by autorepression remain unknown Although *Nanog* autorepression was shown to operate independently of OCT4/SOX2(16) and dependent on ZFP281(13), it is unclear how the NANOG protein levels are perceived by cells to trigger autorepression. Here, we show that ERK signaling is essential for *Nanog* autorepression. NANOG induces *Fgfr2* and *Fgfbp1* exclusively in the high-*Nanog* ESCs to trigger feedback repression by autocrine-mediated activation of ERK signaling. We show that pERK1/2 recruits NONO to the *Nanog* locus to repress *Nanog* transcription by affecting POL2 loading. We show that the Nanog autoregulation process results in a self-perpetuating NANOG-pERK reciprocal regulatory loop. Our results establish that the NANOG-pERK reciprocal regulatory loop is the basis of ERK signaling dynamics and pERK1/2 heterogeneity in pluripotent stem cells. Together with our data show that the NANOG-pERK axis may not merely be viewed as a mechanism to regulate *Nanog*, but also a mechanism by which ERK dynamics and heterogeneity is induced in the pluripotent cells.

## Results

### Residual MEK1/2 activity in the ground state prevents complete derepression of *Nanog*

Transcriptional regulation is the major mechanism regulating *Nanog* heterogeneity, biallelic expression, and autorepression(13). To uncouple the influence of MEK1/2 and GSK3*β* on *Nanog* expression in naïve state of pluripotency, we analyzed the activity of *Nanog* promoter reported by GFP in TβC44Cre6 cell line(1) in combinations of MEK1/2 and GSK3*β* inhibitors. TβC44Cre6 cell line is *Nanog* null ESC in which a Neomycin resistance cassette is knocked in into one allele of *Nanog* and GFP into another allele. *Nanog* expression was derepressed above the basal level (SL) in all treatments. *Nanog* promoter activity was higher in SLPD relative to 2iL (Fig. 1A). To analyze NANOG protein dynamics, we generated a NiRFP2A cell line with both endogenous alleles of Nanog expressing NANOG-IRFP fusion protein (Fig. S1A). Higher NANOGiRFP in SLPD (Fig. 1B), confirmed the highest induction of *Nanog* transcript and protein in SLPD. To dismiss the interference of genetic modifications in the *Nanog* locus on its expression(20); we analyzed its expression in E14Tg2a cells. The *Nanog* transcript (Fig. 1C), transcriptional activity (Fig. 1D), and protein (Fig. 1E) were highest in SLPD, unlike OCT4 protein which changed very little (Fig. 1C-E, Fig. S1B). SLPD and 2iL contain 1 *μ*M PD, higher *Nanog* expression in SLPD indicated inefficient repression of *Nanog* in 2iL/SL2i. We analyzed pERK1/2 to investigate possible modulation of MEK1/2 activity by GSK3*β* (21). The pERK1/2 remained undetectable for up to 4 hrs in SLPD and 2iL/SL2i. It gradually increased in 2iL after 8 hrs but remained undetectable in SLPD (Fig. 1F, Fig. S1C). The pERK1/2 in SLCHIR and 2iL significantly exceeded SL and SLPD respectively by 24 hrs (Fig. 1G, Fig. S1D), suggesting a long-term CHIR treatment enhanced MEK1/2 activity in ESCs. Further, the PD and CHIR dose-responsive experiments confirmed that the pERK1/2 positively correlated with the CHIR concentrations (Fig. 1H, I, Fig. S1E-H). Collectively, our data demonstrate that *Nanog* attains higher expression in MEK1/2 inhibition than in 2iL. GSK3*β* activity negatively modulates MEK1/2 activity and its inhibition by CHIR increases pERK1/2 in 2iL over time.

**Fig. 1.**
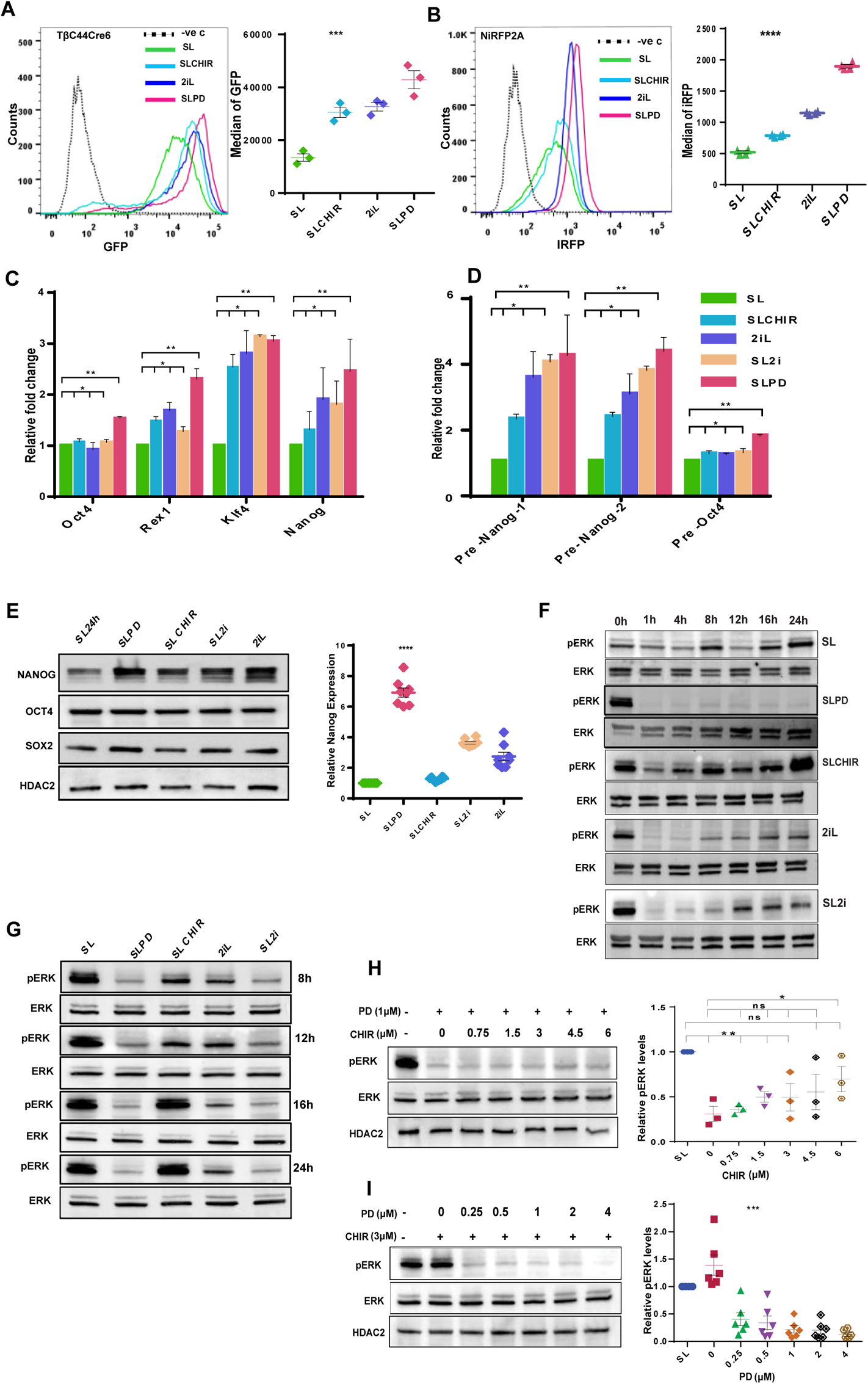
Residual MEK1/2 activity in the ground state prevents complete derepression of *Nanog*. (A) (left) FACS profiles of TβC44Cre6 cultured in indicated conditions for 3 passages. TβC44Cre6 is a *Nanog* null cell line, where *b-geo* cassette is inserted into one allele and GFP into another allele of the *Nanog* gene. The cells were cultured in Serum+LIF (SL) in presence of the 1 *m*M MEK1/2 inhibitor - PD0325901 (SLPD) or 3 *m*M GSK3*b* inhibitor - CHIR99021 (SLCHIR) or in serum-free media - N2B27 with PD0325901, CHIR99021, and LIF (2iL). (right) *Nanog*:GFP population median of TβC44Cre6 (n=3). (B) (Left) FACS profile of NANOG-iRFP protein in NiRFP2A cells cultured in indicated conditions for 3 passages. (right) *NANOG-*iRFP population median of NiRFP2A (n=4). (C) RT-qPCR of pluripotency factors in indicated conditions (SL2i= SL+ PD0325901+CHIR99021). (D) RT-qPCR analysis of pre-mRNA of *Nanog* and *Oct4*. (E) (left)Western blot of NANOG, OCT4, and SOX2. (right) Relative NANOG levels as estimated by densitometry (n=8). NANOG was nearly 7-fold more in PD, which is twice that of 2iL/SL2i. (F) Western blot of pERK1/2 and ERK1/2 at 0, 1, 4, 8, 12, 16, and 24 hrs after media change in indicated treatments. (G) Western blot of pERK1/2 and ERK1/2 in SLPD, SLCHIR, 2iL, and SL2i after 8, 12, 16, and 24 hrs of culture relative to SL, where the cells in SL were harvested 24 hrs after the media change. (H) (left)Western blot of pERK1/2 and ERK1/2 in 1μM PD and increasing concentrations of CHIR in serum-free N2B27 media. (right) Relative pERK1/2 levels (n=3). (I) (left)Western blot of pERK1/2 and ERK1/2 in 3μM CHIR and increasing concentrations of PD in serum-free N2B27 media. (right) Relative pERK1/2 levels (n=6). All error bars in the figure represent s.e.m.

### FGF autocrine signaling pathway components are essential for Nanog autoregulation

We asked if all repressive mechanisms including the *Nanog* autorepression are abolished in SLPD. We generated two NANOG restoration systems by integrating Flag-Avi-NANOG-ER^T2^ (NANOGER^T2^) and a Doxycycline-inducible Flag-Avi-NANOG (FaNANOG) transgene in Tβc44Cre6(1) to derive the TNERT and TDiN cell lines respectively (Fig. 2A, Fig. S2A, B). The repression of endogenous *Nanog*:GFP upon induction of transgenic NANOG by OHT/Dox is a functional readout of *Nanog* autoregulation. *Nanog*:GFP was repressed in OHT/Dox induced TNERT/TDiN in all treatments except SLPD (Fig. 2A, Fig. S2C, D). The data from distinct induction systems conclusively establish an essential role of MEK1/2 in *Nanog* autoregulation.

**Fig. 2.**
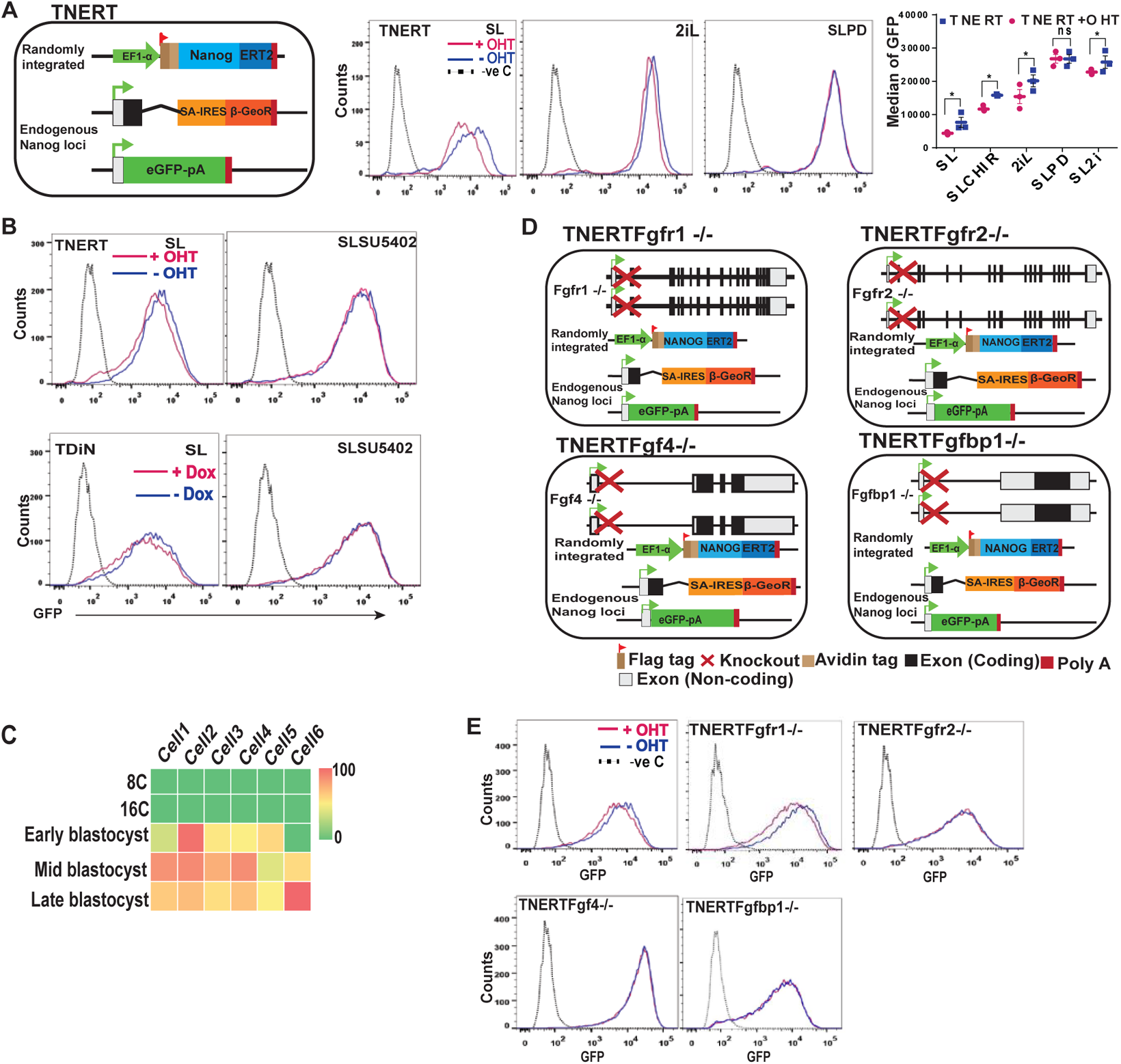
FGF autocrine signaling pathway components are essential for *Nanog* autoregulation. (A) (left) Schematic depiction of Tamoxifen (OHT) inducible TNERT cell line. TNERT and TDiN (Fig. S2A, B) are similar to NERTc3 and 44iN(16), where the NANOG function is reinstated by 4-Hydroxytamoxifen (OHT) or Doxycycline respectively, and endogenous *Nanog* gene activity is reported by GFP. (Middle) FACS profile of TNERT treated with OHT (red) or no OHT (blue). (Right) *Nanog*:GFP population median of TNERT (n=3). (B) FACS profiles of TNERT, and TDiN treated with 2*m*M SU5402, with OHT/Doxycycline (red) or no OHT/Doxycycline (blue). (C) Heat map representing transcript levels (FPKM) of *Fgfbp1* from 8-cell to blastocyst stage analyzed from the single-cell sequencing data. (D) Schematic depiction of TNERTFgfr1−/−, TNERTFgfr2−/, TNERTFgf4−/−, and TNERTFgfbp1−/− cell lines, which are derivatives of TNERT where *Fgfr1*, *Fgfr2*, *Fgf4*, and *Fgfbp1* are knocked out respectively. (E) FACS profiles of TNERT, TNERTFgfr1−/−, TNERTFgfr2−/−, TNERTFgf4−/−, and TNERTFgfbp1−/− cells, treated with OHT (red) or no OHT (blue). All error bars in the figure represent s.e.m.

FGF signaling is the predominant inducer of MEK/ERK in pluripotent cells(22, 23), we investigated its role in autoregulation. NANOGER^T2^/FaNANOG failed to repress *Nanog*:GFP in presence of FGFR inhibitor, suggesting an essential role of FGFRs (Fig. 2B, Fig. S2E). FGFR1, FGFR2, and FGF4 are major receptors and ligands of FGF signaling in early embryos(24, 25). FGFBP1 is a carrier protein expressed from early to late blastocyst (Fig. 2C)(26) that enhances FGF signaling(27). We deleted *Fgfr1*, *Fgfr2*, *Fgf4*, and *Fgfbp1* in TNERT cells to analyze their role in autoregulation (Fig. 2D, Fig. S2F-I). Except in TNERTFgfr1−/−, *Nanog*:GFP was not repressed in TNERTFgfr2−/−, TNERTFgf4−/− and TNERTFgfbp1−/− cells upon OHT induction (Fig. 2E). Our data suggest that FGF autocrine signaling and its components FGFR2, FGF4, and FGFBP1 are essential for *Nanog* autoregulation.

### NANOG enhances the expression of FGFR2, FGF4, and FGFBP1

We analyzed the expression of FGF autocrine signaling components during the time course of OHT induction. *Fgf4, Fgfr2, Fgfr1*, and *Fgfbp1* transcripts were induced within 1-2 hrs (Fig. 3A). Increased pre-mRNA indicated transcriptional activation of these genes (Fig. 3B). ChIP-seq data analysis identified NANOG occupancy on *Fgf4, Fgfbp1, Fgfr1*, and *Fgfr2*, which was further enhanced in *Oct4*+/− cells that have higher NANOG (Fig. S3A)(28). To analyze the dosage-dependent occupancy of NANOG on these genes, we generated EDiN cell line by introducing a Doxycycline-inducible FaNANOG transgene in E14Tg2a. ChIP-PCR confirmed NANOG occupancy on *Fgf4*, *Fgfbp1*, *Fgfr1*, and *Fgfr2*, which was further enhanced in PD (Fig. 3C) and EDiN+Dox (Fig. 3D) which express higher NANOG. The data suggest a dose-dependent occupancy of NANOG on the FGF signaling component genes. FGFR1, FGFR2, and pERK1/2 were significantly increased upon OHT induction in TNERT (Fig. 3E). The strength of FGF signaling depends on facilitation by carrier proteins(27), the affinity of ligands(29, 30) and subsequent subcellular trafficking of the FGFRs(31, 32). The induction of NANOGER^T2^ enhanced FGFR2 on the cell surface (Fig. 3F), unlike the FGFR1 (Fig. S3B) suggesting NANOG specifically enhances FGFR2. Intriguingly, FGFR2 expression exhibited a negatively skewed bimodal distribution resembling *Nanog* expression(1) (Fig. 3F). The NANOGER^T2^ induction increased FGF4 and FGFBP1 secretion by TNERT (Fig. 3G, H, Fig. S3C, D). Collectively the data shows that increased NANOG enhances FGFR2 on the cell surface, and secretion of FGF4 and FGFBP1 to intensify the FGF autocrine signaling. NANOG induces and enhances FGF autocrine signaling through FGFR2 to execute *Nanog* autoregulation.

**Fig. 3.**
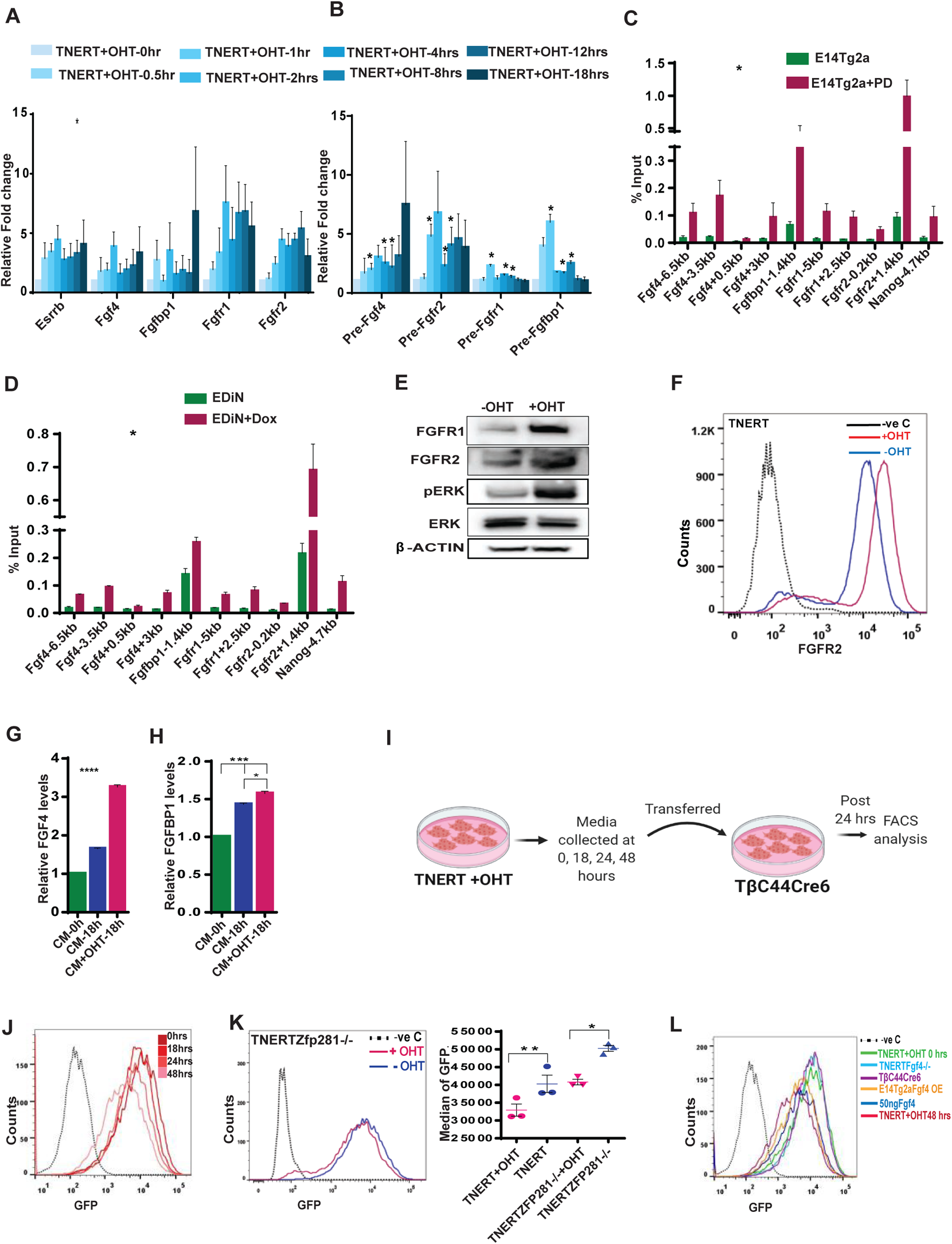
NANOG triggers autoregulation by inducing the expression of FGFR2, FGF4, and FGFBP1. (A) RT-qPCR showing relative transcript levels after 0, 0.5, 1, 2, 4, 8, 12 and 18 hrs OHT treatment in TNERT (n=3). *Esrrb*, a known direct target of NANOG was used as positive control. (B) RT-qPCR of relative levels of pre-mRNA at the above indicated time points after OHT treatment in TNERT (n=3). (C) ChIP analysis of NANOG on *Fgf4*, *Fgfbp1*, *Fgfr1*, *Fgfr2*, and *Nanog* genes in E14Tg2a cells cultured in SL or SLPD for 48 hrs (n=4). (D) ChIP analysis of NANOG on promoters of above-indicated loci in EDiN cells cultured in Doxycycline (red) or no Doxycycline (blue) for 48 hrs (n=3). (E) Western blot of FGFR1, FGFR2, and pERK1/2 in TNERT after 18 hrs treatment with or no OHT. (F) FACS analysis of FGFR2 on the cell surface of TNERT treated with (red) or no OHT (blue) (n=3). (G-H) ELISA-based relative quantification of FGF4 (G) and FGFBP1 (H) in conditioned media from TNERT treated with or no OHT (n=3). (I) Schematic of conditioned media experiment. (J) FACS analysis of Tβc44Cre6 cell line in conditioned media collected from TNERT treated with OHT after 0, 18, 24, and 48 hrs. (K) (left) FACS analysis of TNERTZfp281−/− cells treated with (red) or with no OHT (blue) treatment. (right) *Nanog*:GFP population median of TNERTZfp281−/− (n=3). (L) FACS analysis of Tβc44Cre6 cell line in conditioned media from, TNERT+OHT 0 hrs, TNERTFGF4−/−, Tβc44Cre6 48 hrs, E14Tg2a-FGF4-OE (overexpression) 48 hrs, TNERT+OHT 48 hrs and 50ng/ml FGF4. All error bars in the figure represent s.e.m.

### *Nanog* autoregulation is a cell non-autonomous process mediated by FGF autocrine/paracrine signaling

*Nanog* autorepression is suggested to operate by a cell-autonomous process through intracellular proteins NANOG, ZFP281, and NURD complex(13). Cell non-autonomous function of *Nanog* in the induction of primitive endoderm(33, 34) and essentiality of secreted proteins FGF4 and FGFBP1 in autoregulation prompted us to investigate cell non-autonomous mechanisms. We assessed the ability of conditioned media from OHT induced TNERT cells, to repress *Nanog*:GFP in Tβc44Cre6 lacking *Nanog* (Fig. 3I). The conditioned media was sufficient to repress the *Nanog*:GFP (Fig. 3J, Fig. S3E), suggesting that autoregulation operates via cell non-autonomous mechanisms and discounts the direct role of NANOG in autoregulation as proposed earlier(13). NANOG seems to be essential for triggering autoregulation through FGF autocrine signaling but does not participate in repression. Further, the repression of *Nanog:*GFP in TNERTZfp281−/− cell line lacking Zfp281 (Fig. S3F) suggests that ZFP281 is dispensable for *Nanog* autoregulation (Fig. 3K).

To evaluate if FGF4 secretion was the causative factor of *Nanog* autoregulation in the conditioned media, we treated Tβc44Cre6 with conditioned media from cells with loss or gain of FGF4. The conditioned media from an E14Tg2a cell line overexpressing FGF4 or supplementation of FGF4 (50ng/ml) could repress *Nanog*:GFP. Conversely, the conditioned media from OHT induced TNERTFgf4−/− cells failed to repress *Nanog*:GFP, suggesting FGF4 is the key secreted factor essential for *Nanog* autoregulation (Fig. 3L, Fig. S3G). The ELISA analysis confirmed the secretion and accumulation of FGF4 and FGFBP1 in the conditioned media (Fig. S3H-K). Collectively, our data establish that *Nanog* autoregulation is a cell non-autonomous process triggered by NANOG by augmenting FGF autocrine signaling.

### NANOG induced FGFR2 triggers autoregulation predominately in the ES cell population with higher *Nanog* expression

*Nanog* autoregulation was proposed to restrict NANOG levels within limits to retain the differentiation potential(13, 16). Autoregulation is expected to operate only in *Nanog*-high cells in a population. To evaluate this logic, we used TDiN cell lines with different induction levels of FaNANOG (Fig. S4A). The strength of *Nanog* autoregulation was found to be dependent on the FaNANOG levels and was completely abolished in TDiN clones with low FaNANOG (Fig. S4B, C). Further, the *Nanog:*GFP was repressed only in 10% population of the TNERT with the highest *Nanog* expression but not in the lowest 10% (Fig. 4A). Our experiments conclude that *Nanog* autoregulation predominately operates in a subpopulation of cells with higher *Nanog*.

**Fig. 4.**
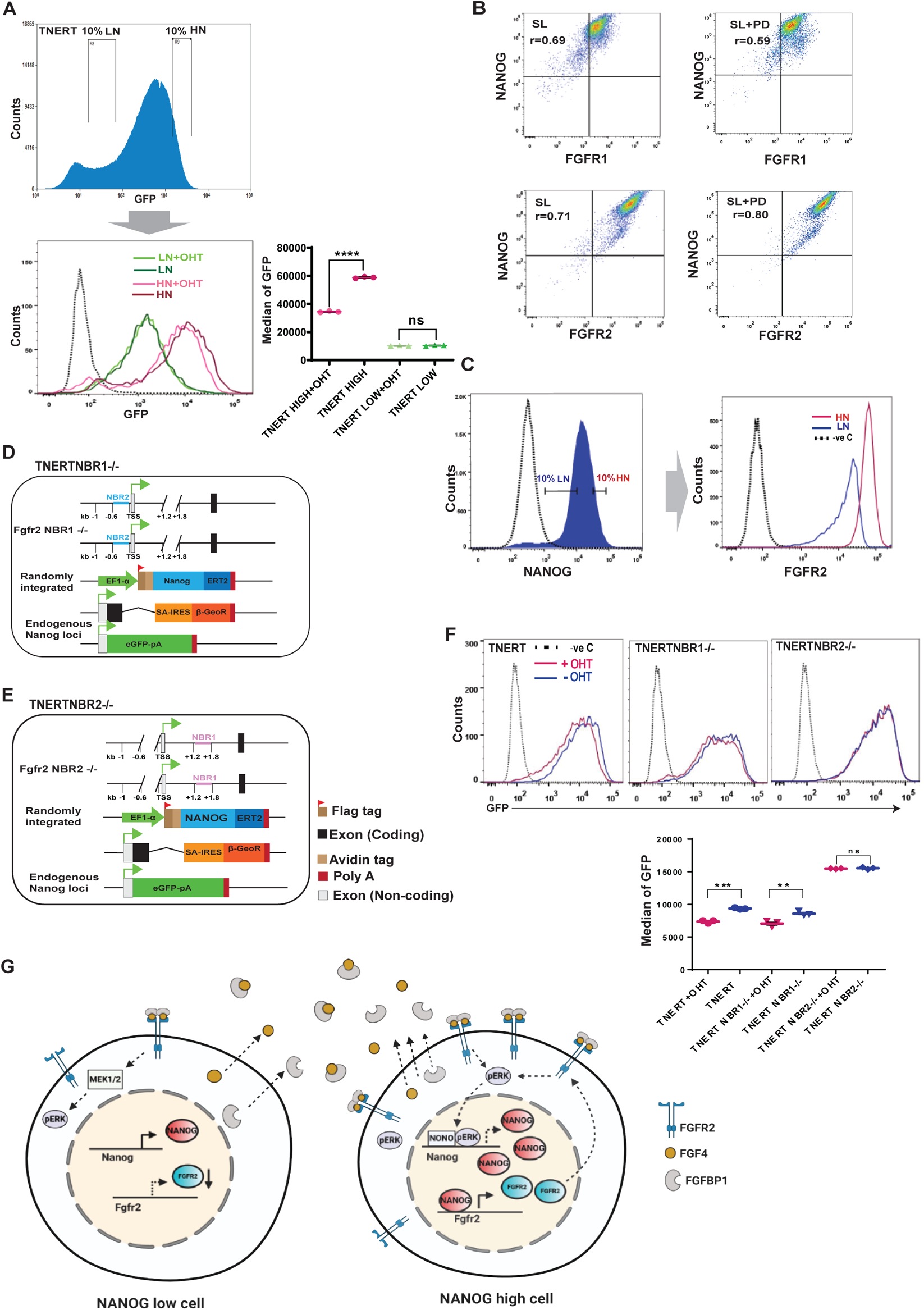
NANOG induced FGFR2 triggers autoregulation predominately in the ES cell population with higher *Nanog* expression. (A) (top) To analyze autoregulation in *Nanog*-high and *Nanog*-low cells, we sorted the lowest and the highest 10% population of the TNERT expressing GFP and treated with OHT. FACS profile of TNERT, the position of the gates indicates the 10% low-*Nanog*:GFP (LN) and 10% high-*Nanog*:GFP (HN) population sorted for culture. (Bottom left) FACS profiles of LN and HN after 18 hrs culture in SL. LN (dark green), HN (dark maroon) in SL, and LN (light green), HN (light maroon) in SL+ OHT. (Bottom right) *Nanog*:GFP population median of TNERT (n=3). (B) FACS profile of E14Tg2a cultured in SL or SLPD for 48 hrs and co-immunostained with anti-NANOG and anti-FGFR1 or anti-FGFR2 antibodies. r-values represent the average of 3 independent experiments (n=3). (C) (left) FACS profile of E14Tg2a immunostained with anti-NANOG and anti-FGFR2 antibody, the gates mark the 10% low-NANOG (LN) and 10% high-NANOG (HN) population. (right) Histogram depicting the FGFR2 expression profiles in the gated LN and HN cell population (n=3). (D-E) Schematic representation of TNERTNBR1−/− (D) and TNERTNBR2−/− (E) cells, in which NANOG binding sequences at +1.4 kb (NBR1) and −0.2 kb (NBR2) are deleted respectively. (F) (Top) FACS profiles of TNERT, TNERTNBR1−/−, and TNERTNBR2−/− with (red) or no OHT treatment (blue). (bottom) *Nanog*:GFP population median of TNERT, TNERTNBR1−/− and TNERTNBR2−/− with or no OHT treatment. (G) A cartoon depicting *Nanog* autoregulation in *Nanog*-high cells. The *Nanog*-high cells secrete more FGF4 and FGFBP1. They contain higher levels of FGFR2 on the surface and are hence more sensitive to the FGF ligand triggering a stronger FGF signaling. The increased pERK1/2 in these cells recruit NONO to the *Nanog* locus and represses *Nanog* transcription. The *Nanog*-low cells secrete very little FGF4 and FGFBP1 and present fewer FGFR2 on their surface and are less sensitive to FGF signaling. The pERK1/2 levels in *Nanog*-low cells are insufficient to execute *Nanog* autoregulation. All error bars in the figure represent s.e.m.

FGF4 and FGFBP1 are secreted proteins, hence cannot distinguish between the *Nanog*-high and low cells in culture. Whereas FGFRs are essential for autoregulation and are retained on the cells, we asked if FGFRs distinguish *Nanog*-high cells from low cells in a population. We analyzed the correlation between the expression of FGFR1, FGFR2, and NANOG in E14Tg2a by FACS. FGFR2 and FGFR1 showed a strong correlation with NANOG, which was further enhanced for FGFR2-NANOG in SLPD (r=0.80) whereas decreased for FGFR1-NANOG (r=0.59) in SLPD (Fig. 4B,) where NANOG levels are higher. These data suggested FGFR2 but not FGFR1 expression levels correlate and respond to NANOG concentration in the cells. FACS analysis showed high FGFR2 in the 10% NANOG high population and lower FGFR2 in the 10% NANOG low population (Fig. 4C). We analyzed the NANOG binding sequences in the *Fgfr2* locus. Two NANOG binding regions (NBR) with multiple NANOG binding sequences were identified in the *Fgfr2* locus from the ChIP-seq(28), NBR1 at 1.4 kb, and NBR2 at −0.2 kb relative to TSS of *Fgfr2*. NBR1 and NBR2 were deleted in TNERT (Fig. 4D, E, Fig. S4D, E). Autoregulation was operational in TNERTNBR1−/− albeit at reduced strength, whereas it was abolished in TNERTNBR2−/− (Fig. 4F), suggesting that NBR2 is essential for the binding of NANOG and activation of *Fgfr2* to trigger autoregulation. Together, our data suggest dose-responsive induction of *Fgf4*, *Fgfbp1*, and *Fgfr2* by NANOG. The *Nanog*-high cells secrete more FGF4 and FGFBP1, also express higher FGFR2 receptors. The FGF4 in presence of FGFBP1 binds to FGFR2 to enhance FGF signaling in *Nanog*-high cells to enhance pERK1/2 and repress *Nanog*. The *Nanog*-low cells express relatively low FGFR2, resulting in weak FGF signaling and the absence of autoregulation (Fig. 4G). We propose that FGFR2 distinguish the *Nanog*-high cells from the low cells to activate ERK-driven autoregulation selectively in *Nanog*-high cells.

### ERK1/2 interacts and recruits NONO to repress *Nanog* transcription

FGF signaling represses *Nanog* transcription(18, 35) and regulates *Nanog* heterogeneity and monoallelic expression(12, 36, 37). How FGF signaling downstream kinases repress *Nanog* is unclear. ERK can induce Tcf15 to repress *Nanog*(38) or it can interact with NONO to regulate bivalent genes(39). We deleted *Tcf15* and *Nono* in TNERT to generate TNERTTcf15−/− and TNERTNono−/− cell lines to examine their function in autoregulation (Fig. 5A, Fig. S5A, B). OHT treatment failed to repress *Nanog*:GFP in TNERTNono−/−, unlike in TNERTTcf15−/− indicating an essential role of NONO but not TCF15 (Fig. 5B, Fig. S5A, C). NONO has been shown to activate ERK1/2(39), and pERK1/2 was substantially reduced in TNERTNono−/− despite OHT induction, unlike in TNERT (Fig. 5C, Fig. S5D). Endogenous immunoprecipitation showed an interaction between NONO and ERK1/2, the interaction was maintained in the presence or absence of NANOG (Fig. 5D). NONO colocalizes with ERK1/2 to bivalent developmental genes to maintain poised POL2(39). The ChIP-seq data analysis from Ma et. al.,(39) and Tee et. al.,(40) showed NONO and ERK1/2 occupancy on the *Nanog* (Fig. S5E). We induced or repressed the pERK1/2 by treatment of E14Tg2a cells with FGF4 or PD (Fig. 5E) and analyzed the occupancy of NONO, pERK1/2, POL2, H3K4me3, and H3K27me3. The transcription start site (TSS) and 5 kb upstream region (−5kb) are the two hubs of transcription factor binding and control of *Nanog* transcription(41, 42). We performed ChiP-qPCR analysis with multiple primer sets spanning the −5.8 kb to +1.5 kb region relative to TSS (Fig. 5F). pERK1/2 and NONO binding was detected in immediate downstream regions of the −5kb, and TSS. Their binding was reduced significantly in PD and enhanced in FGF4 suggesting pERK1/2 and NONO binding on *Nanog* is dependent on FGF signaling (Fig. 5G, H). pERK1/2 was shown to recruit NONO to bivalent genes(39). Although *Nanog* is not a bivalent gene, our data suggests pERK1/2 recruits NONO to *Nanog*. POL2 occupancy seen in TSS and downstream region was reduced in FGF4 and enhanced in PD treatment suggesting active transcription of *Nanog* in PD and repression in FGF4 (Fig. 5I). This was corroborated with enhanced enrichment of the transcription activating histone mark H3K4me3 in PD (Fig. 5J) and enrichment of transcription repressive mark H3K27me3 at the −5kb region of the *Nanog* in FGF4 treatment (Fig. 5K). pERK1/2 phosphorylates NANOG, USP21 and affects NANOG stability and transactivation capability(19, 43, 44). In agreement with NANOG destabilization by pERK1/2(19, 43, 45), The half-life of NANOG was significantly compromised in FGF4 treated cells but enhanced in PD (Fig. 5L, Fig. S5F), suggesting that the FGF/ERK represses *Nanog* transcription and also affects NANOG stability. Collectively, these data suggest that FGF signaling activates pERK1/2 and its binding onto *Nanog* in a concentration-dependent manner. pERK1/2 is essential for the recruitment of NONO to the *Nanog* locus. pERK1/2-NONO are known to poise the POL2 in bivalent genes(39). In contrast, pERK1/2-NONO affects POL2 loading onto the *Nanog* locus preventing the initiation of transcription. In the absence of active FGF signaling, pERK1/2-NONO occupancy on the *Nanog* is decreased permitting increased POL2 loading and transcription activation of the *Nanog* (Fig. 5M).

**Fig. 5.**
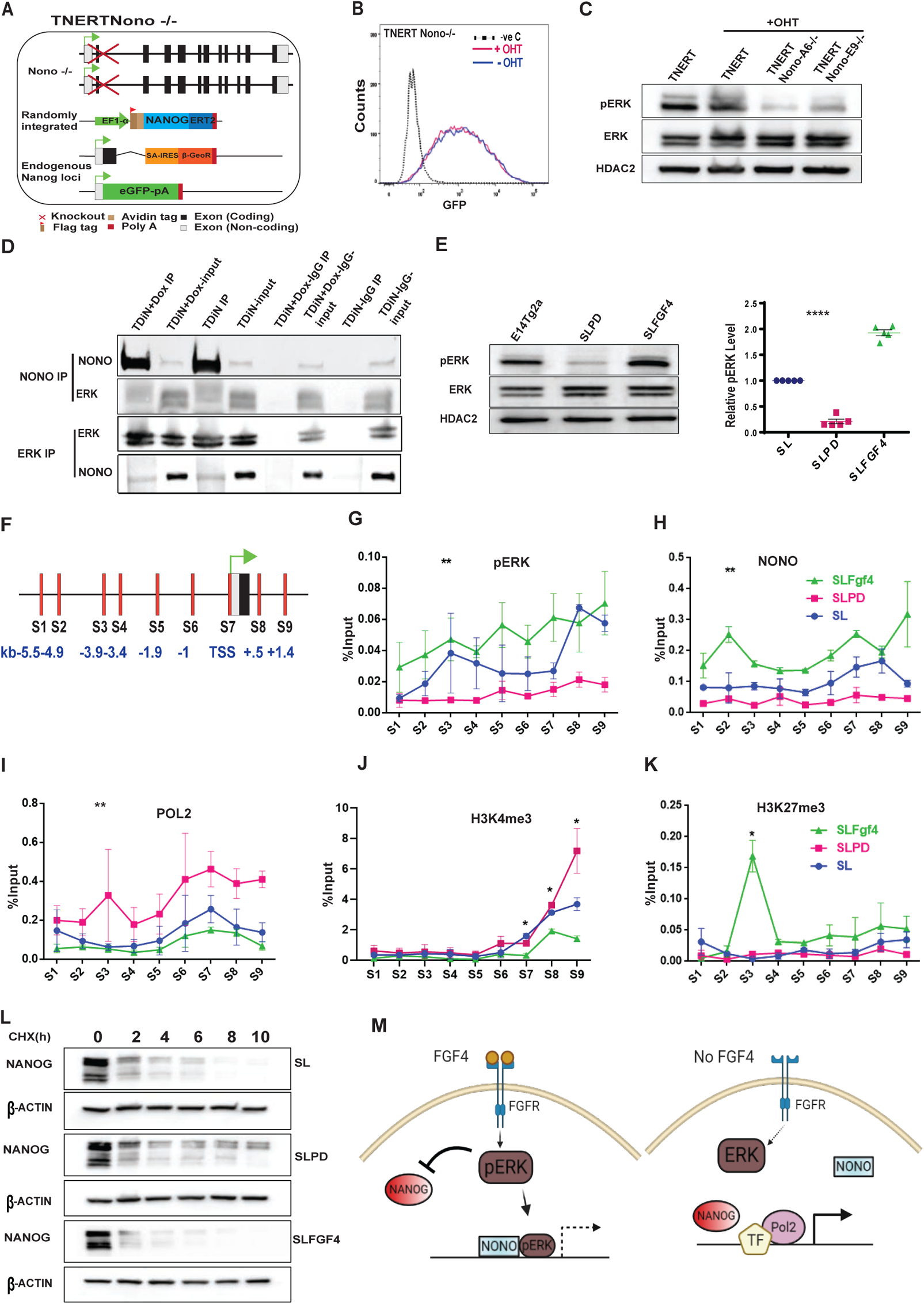
ERK1/2 interacts and recruits NONO to repress *Nanog* transcription. (A) Schematic of TNERTNono−/− cell line; a derivative of TNERT in which *Nono* is knocked-out. (B) FACS profile of TNERTNono−/− treated with or no OHT (n=3). (C) Western blot of pERK1/2 and ERK1/2 in TNERT and TNERTNono−/− cells treated with or no OHT (n=3). (D) Immunoprecipitation analysis showing interactions between ERK1/2 and NONO in the presence or absence of *Nanog* induction by Doxycycline in TDiN cells. (E) (left) Western blot of pERK1/2 and ERK1/2 in E14Tg2a cells treated with PD or FGF4. (right) Relative levels of pERK1/2 in E14Tg2a cells treated with PD or FGF4 (n=4). (F) Schematic representation of *Nanog* locus comprising the −6.0 to +2kb region. The vertical bars represent relative positions of primer pairs used for ChiP-qPCR analysis. S1-S6 are located upstream of the TSS, S7 primer pair is located around TSS, S8 and S9 are located downstream in the first intron. (G-K) ChIP-qPCR analysis of pERK1/2 (G), NONO (H), Pol2 (I), H3K4me3 (J) and H3K27me3 (K) on *Nanog* 5’ region in E14Tg2a cells (blue), treated with FGF4 (green) and with PD (pink) (n=3). (L) Cycloheximide chase assay of NANOG in SL, SLPD, and SLFGF4 in E14Tg2a cells. (M) A cartoon illustrating the repression of *Nanog* by FGF signaling and derepression of *Nanog* in absence of FGF signaling. The FGF4 activates the FGF signaling cascade, resulting in phosphorylation of ERK1/2. pERK1/2 interacts and recruits NONO to the *Nanog* promoter and represses transcription of *Nanog*. pERK1/2 also affects the stability of the NANOG. In absence of FGF4, the pERK1/2 levels decrease resulting in enhanced stability of NANOG and transcription of *Nanog* locus by NANOG and other pluripotency factors resulting in derepression of *Nanog* locus. All error bars represent s.e.m.

### NANOG regulates ERK signaling dynamics and heterogeneity

ERK signaling regulates *Nanog* expression and heterogeneity in ES cells. Recently pERK1/2 expression is reported to be heterogeneous and dynamic in ES cells and the preimplantation embryos(24, 25, 46). We have shown that FGFR2 exhibits a negatively skewed bimodal expression similar to *Nanog* in ESCs (Fig. 3F) and *Fgfr2* is induced in a dosage-dependent manner by NANOG. We asked if NANOG dynamics could regulate ERK signaling dynamics in ES cells through *Fgfr2*. Immunostaining showed heterogeneous expression of NANOG and pERK1/2 in WT ESCs (E14Tg2a) with some cells co-expressing both (Fig. 6A). Their expression showed a strong correlation (r=0.6675) suggesting a positive association between NANOG and pERK1/2; similar to NANOG and FGFR2. NANOG showed a broad range of expression as represented by the broad range of relative fluorescence intensity (RFI 0-20000), pERK1/2 showed a relatively narrow range of expression (RFI 1500-3000) in ESCs (Fig. 6B). The pERK1/2 expression in *Nanog* null ESC (TBC44Cre6) was very low relative to WT ESCs (RFI <2000), suggesting pERK1/2 levels are dependent on NANOG. NANOG overexpression in WT ESCs (EDiN) enhanced pERK1/2 levels by multiple folds relative to WT and broadened the range of pERK1/2 levels (RFI >30000) (Fig. 6A) with a moderate correlation between NANOG and pERK1/2 (r = 0.5375) (Fig. 6B). Intriguingly, high levels of pERK1/2 failed to repress *Nanog* transgene and significantly reduce NANOG in EDiN. This resulted in the coexistence of high pERK1/2 and high NANOG in the cells. Despite very high levels of pERK1/2, *Nanog* over-expressing EDiN does not differentiate suggesting that the *Nanog* function in ESC self-renewal is dominant over the pERK1/2 function in the differentiation of ESCs. These data suggest that pERK1/2 expression levels and dynamic range of expression in ESCs are dependent on the expression level of *Nanog* and its dynamics. To further validate this, we isolated *Nanog*-high subpopulation cells by sorting the highest 10% iRFP expressing NiRPF2A reporter ESCs by FACS. The expression of pERK1/2 and NANOG was analyzed in these cells every 4 hours during their culture to study the dynamics of NANOG and pERK expression. The sorted cells expressed NANOG and pERK1/2. After 4 hours of culture in fresh media, the NANOG expression increased with a concomitant decrease in pERK1/2 (NANOG high-pERK1/2 low state). After 8 hours, the NANOG expression decreased and the pERK1/2 expression increased. At 12 hrs the cells showed relatively low pERK1/2 and low NANOG expression (Fig. 6C). A relative median fluorescence intensity plot of NANOG and pERK1/2 suggests that NANOG and pERK1/2 follow a dynamic cycle of expression during culture (Fig. 6D). This was further confirmed by immunostaining and imaging of the sorted cell line at every 4 hr intervals (Fig. S6A). These results suggest that ESCs continuously transit between different states of NANOG and pERK1/2 expression resulting in heterogeneous and dynamic expression pERK1/2.

**Fig. 6.**
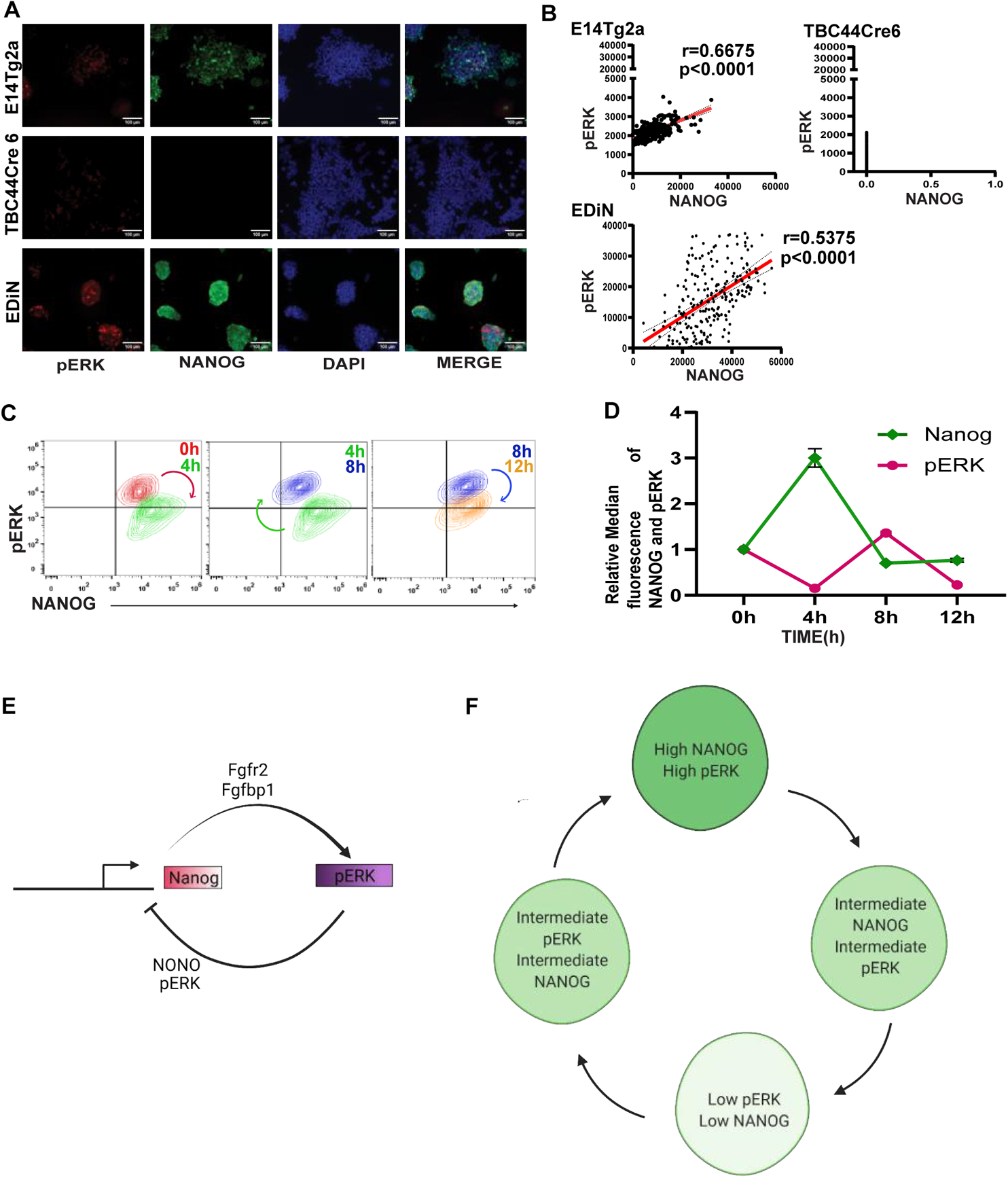
NANOG regulates ERK signaling dynamics and heterogeneity. (A) Immunofluorescence of pERK1/2 (red) and NANOG (green) in the indicated ESCs. (B)The normalized fluorescence intensity of pERK1/2 was plotted against the normalized fluorescent intensity of NANOG. (C) Contour plot of FACS analysis of pERK1/2 and NANOG in 10% NANOG-high NiRFP2A cells cultured for the indicated time. (D) A plot of median fluorescence intensity of pERK1/2 and NANOG relative to 0 hrs culture of 10% NANOG-high NiRFP2A cells. The NANOG and pERK1/2 expression oscillate between high and low levels in the cells during the course of culture. (E) A working model of the NANOG-pERK1/2 reciprocal regulatory loop operating in ESCs. NANOG induces *Fgfbp1* and *Fgfr2 to* enhance ERK signaling in *Nanog*-high cells. pERK1/2 along with NONO occupy the *Nanog* promoter to repress its transcription. The transcription repression results in reduced NANOG, which prevents induction of *Fgfbp1*and *Fgfr2.* The is reduces ERK activity relieving the repression on the *Nanog* promoter. (F) A schematic depicting the progression of cells through different expression states of NANOG and pERK1/2 expression in the ESC population. The cells expressing high-NANOG induce *Fgfbp1* and *Fgfr2* to activate pERK by autocrine signaling to give rise to a high-NANOG:high-pERK state. The repression of *Nanog* transcription by pERK leads the cells through different intermediate levels of expression of NANOG and pERK resulting in a low-NANOG:low-pERK state. The low pERK permits transcription of *Nanog* and gradual induction of *Fgfbp1* and *Fgfr2* by NANOG culminating in a high-NANOG:high-pERK state. The cells will cycle through different levels of pERK and NANOG levels generating a heterogeneous population with a strong correlation between pERK and NANOG in an ESC cells culture.

## Discussion

We demonstrate that the highest possible expression of *Nanog* could be achieved in SLPD by attaining a consistent low MEK1/2 activity. Wnt signaling can activate MEK1/2 at multiple levels(21), a relatively lower level of *Nanog* expression in SL2i and 2iL could be attributed to a time-dependent increase in MEK1/2 activity in presence of PD and CHIR (Fig. 1F, G). The inhibition of MEK1/2 prevents differentiation in 2iL, but a time-dependent increase in MEK1/2 activity is significant and sufficient to facilitate *Nanog* autoregulation. A time-dependent variation in MEK1/2 activity in 2iL opens up the plausibility of other molecular processes regulated by MEK1/2 activity to be functional in a naïve state.

Overexpression of *Nanog* is limited by an autorepression mechanism operating at the transcriptional level to retain the differentiation potential of ESCs(13, 16). Among the multiple possible pathways that can regulate *Nanog*(*17, 35, 38, 44*), we show that FGF autocrine signaling is recruited for *Nanog* autoregulation. We show that a NANOG dosage-dependent differential induction of *Fgfr2* in *Nanog*-high ESCs triggers autoregulation by activation of ERK1/2. pERK1/2 recruits NONO to the *Nanog* locus and affects the loading of POL2 onto the *Nanog* locus reducing *Nanog* transcription (Fig. 5M). Other reports(19, 43) and our data show that pERK1/2 can affect NANOG stability and may contribute to autorepression. However, the inability of pERK1/2 to significantly repress NANOG expressed from a transgene and a strong correlation between NANOG-pERK1/2 (Fig. 6A, B) dismisses the possibility of significant contribution from post-transcriptional mechanisms in autoregulation. Our data suggest that *Nanog* autoregulation is triggered above a threshold of NANOG, thereafter the intensity of repression is dependent on the level of NANOG in the cell.

We show that NANOG activates ERK signaling by inducing *Fgfr2*, *Fgf4*, and *Fgfbp1.* The activated ERK1/2 together with NONO represses transcription of *Nanog*, resulting in a NANOG-pERK1/2 reciprocal regulatory loop (Fig. 6E). The subpopulation of ES cells expressing high NANOG will have higher FGFR2. This induces high ERK activity resulting in a high-NANOG:high-pERK state. The repression of *Nanog* transcription by pERK in these cells reduces NANOG, reducing transcription of *Fgfr2*. The cells traverse through various intermediate levels of NANOG and pERK1/2 resulting in a low-NANOG:low-pERK state. Low pERK1/2 permits activation of NANOG by other pluripotency factors gradually increasing NANOG in these cells. The increased NANOG activates *Fgfr2*, *Fgfbp1*, and *Fgf4* to induce ERK activity leading to various intermediate levels of NANOG and pERK culminating in high-NANOG:high-pERK state. This induces a self-perpetuating cycle of activation of ERK signaling by NANOG and repression of *Nanog* by pERK1/2 leading to dynamic expression levels of NANOG and pERK1/2 in the ESC population (Fig. 6F).

pERK1/2 heterogeneity is suggested to be a vital determinant of fate choice in ICM and ES cells(24, 25, 45, 47, 48). The mechanism generating pERK1/2 heterogeneity is unclear. pERK1/2 heterogeneity may originate due to differential local concentrations of FGF4 or FGFBP1 or heterogeneous expression of receptors FGFRs or by negative feedback regulators (ETV5, DUSP1/6). *Nanog* is considered to induce FGF paracrine signaling through FGF4 secretion and specify primitive endoderm by cell-autonomous mechanisms(33, 34). Although FGF4 is essential for *Nanog* autoregulation, it is a secreted protein. Its induction by NANOG can neither explain the functioning of autoregulation exclusively in *Nanog*-high cells nor the heterogenous pERK1/2 activation in ESCs or ICM. FGFR1 is unlikely to induce ERK1/2 heterogeneity as it is relatively uniformly expressed in the epiblast(24, 25) and ESCs. Dosage-dependent induction of *Fgfr2* by NANOG and its accumulation on the surface of NANOG high cells can potentiate the cells to differentially respond to FGF4. Our data establish that the dosage-dependent induction of *Fgfr2* is the basis for differential activation of ERK1/2 in subpopulations of ESCs resulting in pERK1/2 heterogeneity. The carrier protein FGFBP1 may also locally enhance FGF signaling further contributing to pERK1/2 heterogeneity similar to heparan sulfate proteoglycans(49).

We propose the reciprocal regulation of *Nanog* by ERK signaling and ERK signaling by NANOG as the basis for both NANOG and pERK1/2 heterogeneity. We suggest that the NANOG-pERK axis may not merely be viewed as a mechanism of regulation of *Nanog* expression by ERK signaling, rather as a cyclic circuit where *Nanog* heterogeneity and expression dynamics lead to ERK signaling dynamics and vice versa. *Nanog* and ERK signaling are induced in multiple cancers(50, 51). The significance of the NANOG-pERK1/2 reciprocal regulatory loop in establishing heterogeneity and ERK signaling dynamics may not be limited to pluripotent cells but could be relevant in cancer stem cells and tumor heterogeneity.

## Methods

### Cell Culture

The cell lines used in this study and their origin is depicted in Fig. S7. All the cells used in this study are derivatives of E14Tg2a ES. The cells were cultured as described earlier (2). 4-Hydroxytamoxifen (4-OHT), Doxycycline, and Cycloheximide were used at a concentration of 1 μg/ml, 1 μg/ml, and 100 μg/ml respectively. The TNERT and its derivative cell lines were treated with 4-OHT for 18 hrs except when indicated. TDiN and EDiN were treated with Doxycycline for 48 hrs unless indicated. CHIR99021 (CHIR, PD0325901 (PD), and SU5402 were used at 3 μM, 1 μM, and 2 μM, respectively, except when indicated. FGF4 and FGFBP1 were used at 50ng/ml concentration. The cells were cultured in Serum+ LIF (SL), SL+ PD (SLPD), SL+CHIR (SLCHIR), SL+SU5402 (SLSU5402), SL + PD +CHIR (SL2i) and N2B27+LIF+PD+CHIR (2iL) for at least 2 passages before treating with either 4-OHT or Doxycycline.

The cells were cultured on cell culture dishes coated with 0.1% gelatin for all experiments. The conditioned media from the cells was collected after the specific treatments or indicated time points. The conditioned media was passed through a 0.22 μM filter and added to Tβc44Cre6 or TNERT cells. The cells were cultured in the conditioned media for 24 hrs before FACS analysis.

### Generation of Knock-out cell lines using paired CRISPR constructs

pU6-iRFP (pU6-Cas9-T2A-iRFP-2A-PuroR) construct was engineered by replacing mCherry coding sequence with iRFP670-2A-PuroR cassette in pU6-(BbsI)-CBh-Cas9-T2A-mCherry plasmid (Addgene 64324) by Gibson assembly. For generating knock-out of a gene, two sgRNAs were designed with the expected cutting sites at least 30 bps apart to achieve deletion of at least 30 bps or more. For genotyping of the deletions, a set of genotyping primers was designed outside the deletion region flanking the sgRNA pair. The sgRNAs were designed using the UCSC genome browser and Deskgen or Benchling. The sequences of the sgRNAs and the genotyping primers are detailed in Table S1. All sgRNAs were cloned into pU6-Cas9-T2A-iRFP-2A-PuroR plasmids. To generate a paired sgRNA construct, the U6-SgRNA cassette from one plasmid containing the sgRNA was amplified and Gibson assembled into the XbaI site of the plasmid containing the other sgRNA pair of the pair. Around 1 µg of paired sgRNA CRISPR plasmid was nucleofected in 1 million cells. The transfected cells were sorted by FACS for iRFP expression and cultured to obtain clones. The clones were genotyped by PCR using respective primer sets to identify the heterozygous and homozygous clones. The sequence of the derivation of cell lines is described in Fig. S7.

### Generation of Knock-in cell lines

A sgRNA encompassing the stop codon of *Nanog* was cloned into pU6-iRFP and co-transfected with the targeting vectors. The 2A-mCherry cassette was replaced with iRFP sequences by Gibson assembly in Nanog-2A-mCherry targeting vector (Addgene 59995) to generate Nanog iRFP670 fusion targeting vector. Around 3 µg plasmid (targeting vector and CRISPR plasmid) were nucleofected in 3 million E14Tg2a cells. The cells were selected against G418. The derivation of cell lines is described in Fig. S7.

### Real-time PCR analysis

The RNA was extracted with TRIZOL reagent and quantified using a Nanodrop2000 spectrophotometer (Thermo Fisher Scientific). One microgram of total RNA was reversed transcribed into cDNA by using superscript III. All real-time PCR was carried out with Power SYBR Green PCR master mix on the ABI prism 7900 HT sequence detection system (ABI) as per the manufacturer’s instructions. GAPDH was used as an internal control or normalizer. The data was analyzed by SDS 2.2 software provided with the instrument. The primers used for real-time PCR are given in Table S1.

### Western blot analysis

The cells were harvested by using RIPA buffer with 25mM Tris HCl (pH 8.0), 150mM NaCl, 1%NP-40, 0.5% Sodium deoxycholate, 0.1% SDS and Complete Protease Inhibitor Cocktail Tablets (Roche). The protein samples were resolved by 4-20% gradient SDS-PAGE and electroblotted on to polyvinylidene difluoride (PVDF) membrane. The blot was blocked with 3% Blotto for an hour and incubated overnight with a primary antibody at 4°C. Blots were washed thrice with TBST and hybridized with secondary antibody and the blots were visualized using enhanced chemiluminescence (ECL)detection kit. Western blot quantifications were performed using Image lab (Bio-rad).

### Chromatin Immunoprecipitation (ChIP)

Cells were fixed by adding 270 µL of 37% formaldehyde into 10 ml of media and incubated for 10 minutes at 37°C to crosslink the chromatin. Cells were washed twice with cold PBS containing protease inhibitors. Cells were scraped and harvested by centrifugation. The cell pellet was dissolved in 200 µL of SDS Lysis Buffer (1% SDS, 10 mM EDTA, and 50 mM Tris, pH 8.0) containing protease inhibitors (per 10^6^ cells) and incubated on ice for 10 min. The 25 cycles of sonication were used to shear DNA between 200 to1000 base pairs. The sample was centrifuged at 13,000 rpm for 10 min (at 4°C). The supernatant was diluted by adding 1800 µl ChIP Dilution Buffer (1.1% Triton X-100, 1.2 mM EDTA, 16.7 mM Tris-HCl, pH 8.0, 167 mM NaCl with protease inhibitors). The 1% input was aliquoted from the supernatant. To reduce nonspecific background, diluted cell supernatant was preabsorbed for one hour at 4°C with protein A/G magnetic beads (Invitrogen). The supernatant fraction was incubated overnight at 4°C with an appropriate antibody and protein A/G magnetic beads were blocked with 4% BSA, 2µg salmon sperm DNA. The next day, pre-blocked beads were mixed with the sample and incubated for 1 hour to capture the antibodies. The supernatant was discarded and washed in the given order with 1 mL of each of the buffers - Low Salt Wash Buffer (0.1% SDS, 1% Triton X-100, 2 mM EDTA, 20 mM Tris-HCl, pH 8.0, 150 mM NaCl), High Salt Wash Buffer (0.1% SDS, 1% Triton X-100, 2 mM EDTA, 20 mM Tris-HCl, pH 8.0, 500 mM NaCl), LiCl Wash Buffer (0.25 M LiCl, 1% IGEPAL-CA630, 1% deoxycholic acid, 1 mM EDTA, 10 mM Tris, pH 8.0.), and TE buffer. DNA was eluted with elution buffer (1%SDS, 0.1M NaHCO3). The sample input and the ChIP chromatin were reverse crosslinked with 20 µL of 5 M NaCl by heating at 65°C for 4 hours. Followed by one hour at 45°C with 10 µL of 0.5 M EDTA, 20 µL 1 M Tris-HCl, pH 6.5, and 2 µL of 10 mg/mL Proteinase K. Finally, DNA was eluted in 50 µL water using a minEleute PCR purification kit. Then 1µL of sample and input was used for qPCR analysis. The primers used for qPCR analysis were listed in Table S1.

### Co-Immunoprecipitation in ES cells

10-12 million ES cells were harvested by trypsinization, washed twice with cold PBS, and resuspended in 800 µl of CoIP Lysis Buffer (50 mM Tris-HCl, pH 67.5; 350 mM NaCl, 0.7% NP40, EDTA 0.1mM, 20% (v/v) glycerol, and protease inhibitor cocktail). The cell lysate was mixed with protein A/G magnetic beads for 1 hour at 4°C for pre-clearing the background. Then 5% input was aliquoted and the remaining supernatant was incubated overnight with appropriate primary antibody. The protein A/G magnetic beads were blocked overnight at 4°C with 200 µl of CoIP Lysis buffer containing 4% BSA. The next day, the beads were transferred to the primary antibody incubated tubes and incubated for one hour at 4°C. The bead was washed three times with ice-cold TBS150 (50mM Tris, 150mM NaCl) and the protein was eluted with 2X sample buffer (125 mM Tris-HCl, pH 6.8, 4% SDS, 20% (v/v) glycerol, 0.004% bromophenol blue), by boiling for 5 min. The western was done for sample and input and the interaction was analyzed.

### Immunocytochemistry

The cells were cultured in 24 well dishes and fixed in 3.7% formaldehyde diluted in PBS for 15 mins at RT. After 3 washes with PBS, the cells were permeabilized and blocked with PBS containing 0.5% BSA and 0.3% Triton-X100 for 1 hour at room temperature. The cells were hybridized with primary antibody (1:100 dilution) in PBS containing 0.5% BSA at 4°C overnight in a humidified chamber. The cells were washed three times with PBS and hybridized to appropriate secondary antibody at 1:1000 dilution room temperature for 1hour. The nuclei were stained with DAPI in 1X PBS for 20 min at room temperature. The cells were washed thrice with PBS. The cells were layered with 100 µl of the mixture of PBS and Glycerol (1:1) and the images were acquired on the ZEISS Axio observer microscope and analyzed using ImageJ software.

### ELISA Assay

The condition media from the cell lines was collected at the respective time points. 100 µl of the media was coated per well of 96 wells of ELISA plate by incubating overnight at 4°C. The wells were washed thrice with PBS containing 0.05% Tween-20 and blocked with PBS containing 2% BSA for one hour at room temperature. The wells were washed once with PBS and incubated with the appropriate primary antibody (1:100) for one hour. Washed thrice with PBST, an appropriate HRP-labeled secondary antibody was hybridized for one hour at room temperature. The wells were washed thrice with PBST and incubated in substrate solution OPD (o-phenylenediamine dihydrochloride) 3mg/ml with 6 µl/ml H_2_O_2_) for 30 min in dark. The reaction was stopped by using 2N H_2_SO_4_. The absorbance was measured at 492 nm in Power wave XS2 (Bio Tek instruments).

### FACS analysis

#### Reporter cells

Cells were trypsinized and collected by spinning at 800 rpm for 5 min. The media was removed and cells were resuspended in 300 µl of PBS containing 2% FBS at 10^6^ cells/ml. The samples were analyzed in the Gallios flow cytometer (Beckman Coulter) or Fortessa flow cytometer (BD Biosciences). Sorting was performed on a MoFlo-XDP cell sorter (Beckman Coulter).

#### Immunostained cells

Cells were harvested by treatment with 0.5 mM EDTA and resuspended into single cells. The cells were fixed in PBS with 4% paraformaldehyde (PFA) for 20 min at room temperature. Cells were washed twice with cold PBS and incubated with methanol for 30 min for permeabilization. In the case of experiments involving the analysis of FGFRs on the cell surface, the permeabilization step was excluded. Then cells were blocked with PBS containing 0.5% BSA for 60 min at room temperature. The cells were washed and hybridized to the appropriate primary antibody at 4°C overnight. The cells were washed thrice with PBS and hybridized to the appropriate secondary antibody in PBS containing 0.5% BSA at1:1000 dilution for one hour at room temperature. The cells were washed thrice with PBS and the fluorescence profiles were acquired in the Gallios FACS analyzer (Beckman Coulter). All the FACS data were analyzed using FlowJo software (BD Biosciences).

#### Statistical analysis and reproducibility

Statistical analysis was done by using a two-tailed paired or unpaired student t-test. The representation of data is in the form of means+/−SEM. The mean was calculated for more than three independent experiments *P* value<0.05 is considered as statistically significant. * represents P<0.05, ** represents P<0.001, *** represents P<0.0001, and **** represents P<0.00001.

## Acknowledgments

HTK was supported by a fellowship from ICMR (India), DJ, VVV, MS was supported by a fellowship from UGC (India). RSR was supported by DBT grant No: BT/PR14064/GET/119/16/2015. PCS was supported by WT/DBT IA grant No: 500053/Z/09/Z. We thank the Microscopy, and FACS core facilities of CCMB for the support extended to carry out this work. We thank Ian chambers for his constructive comments on the manuscript.

## Author contributions

Conceptualization, H.T.K, and P.C.S Methodology, H.T.K and P.C.S; Investigation, H.T.K, R.S.R., V.V.V., and G.S.; Writing – Original Draft, H.T.K. and P.C.S.; Writing – Review & Editing, H.T.K., and P.C.S.; Funding Acquisition, P.C.S.; Resources, H.T.K, R.S.R., M.S., and D.J.; Visualization, H.T.K, and D.J.; Supervision, P.C.S.

## Competing interest

The authors declare no competing interests.

## Supplementary Information Text

### Subhead. Materials

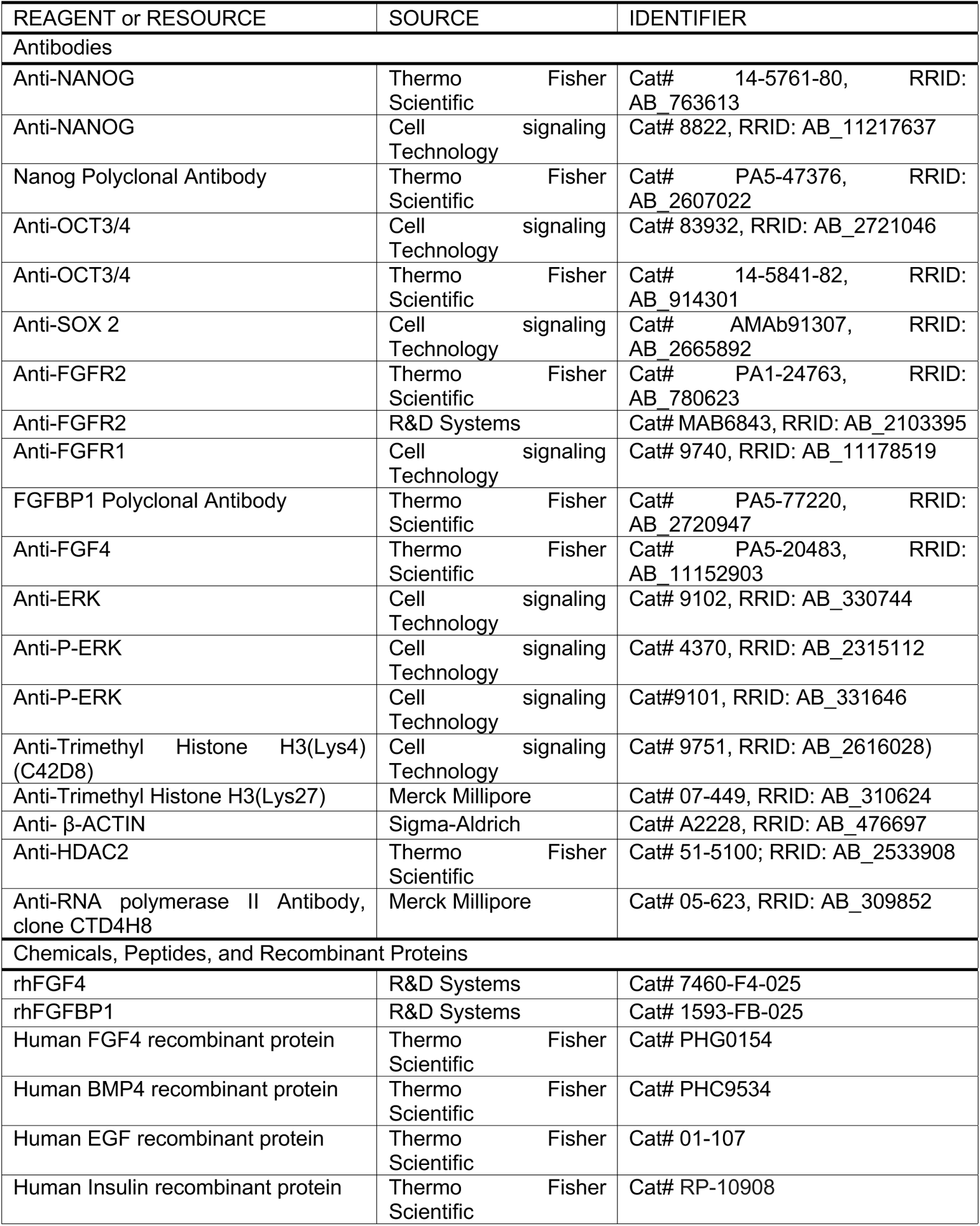
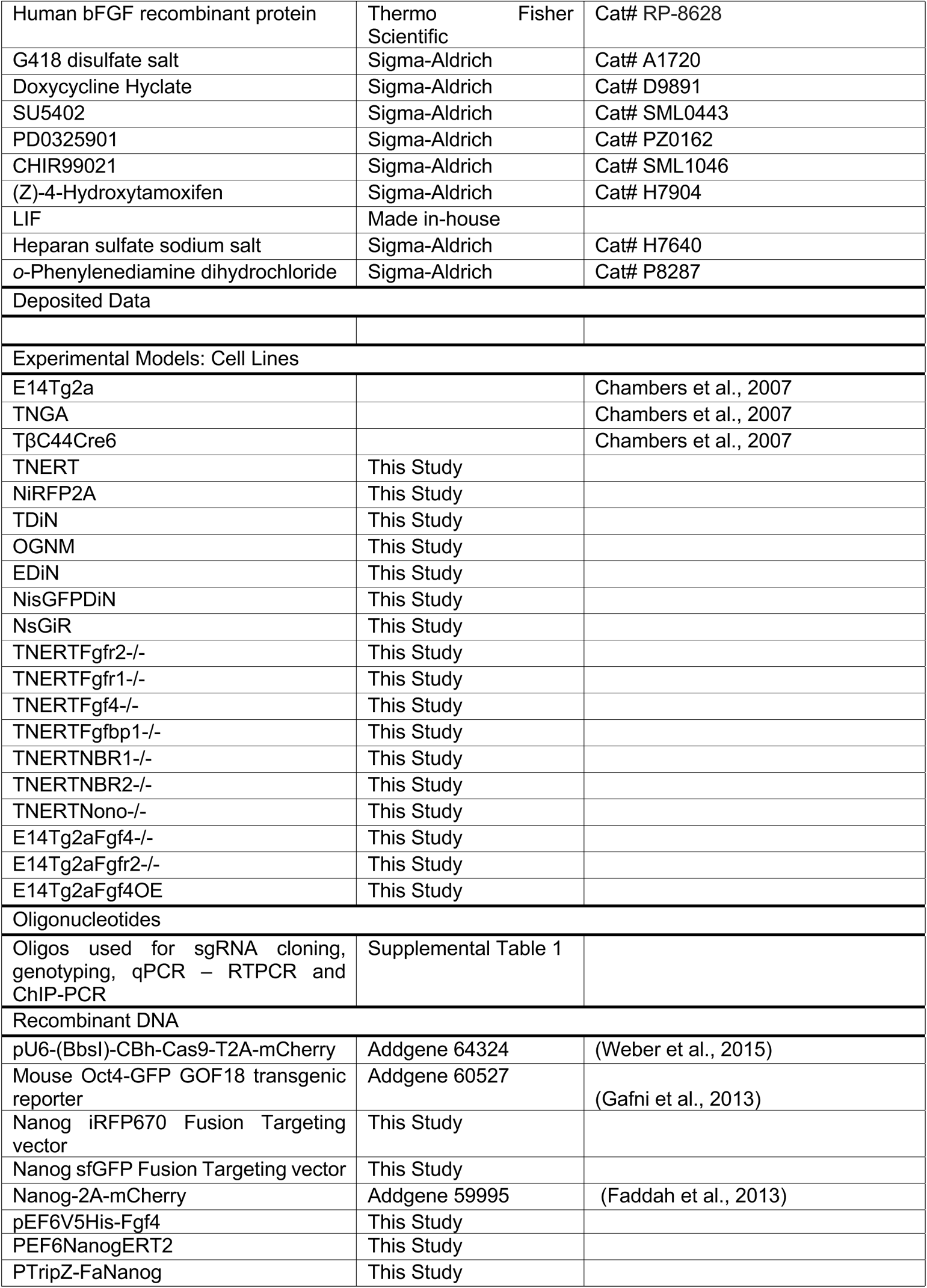
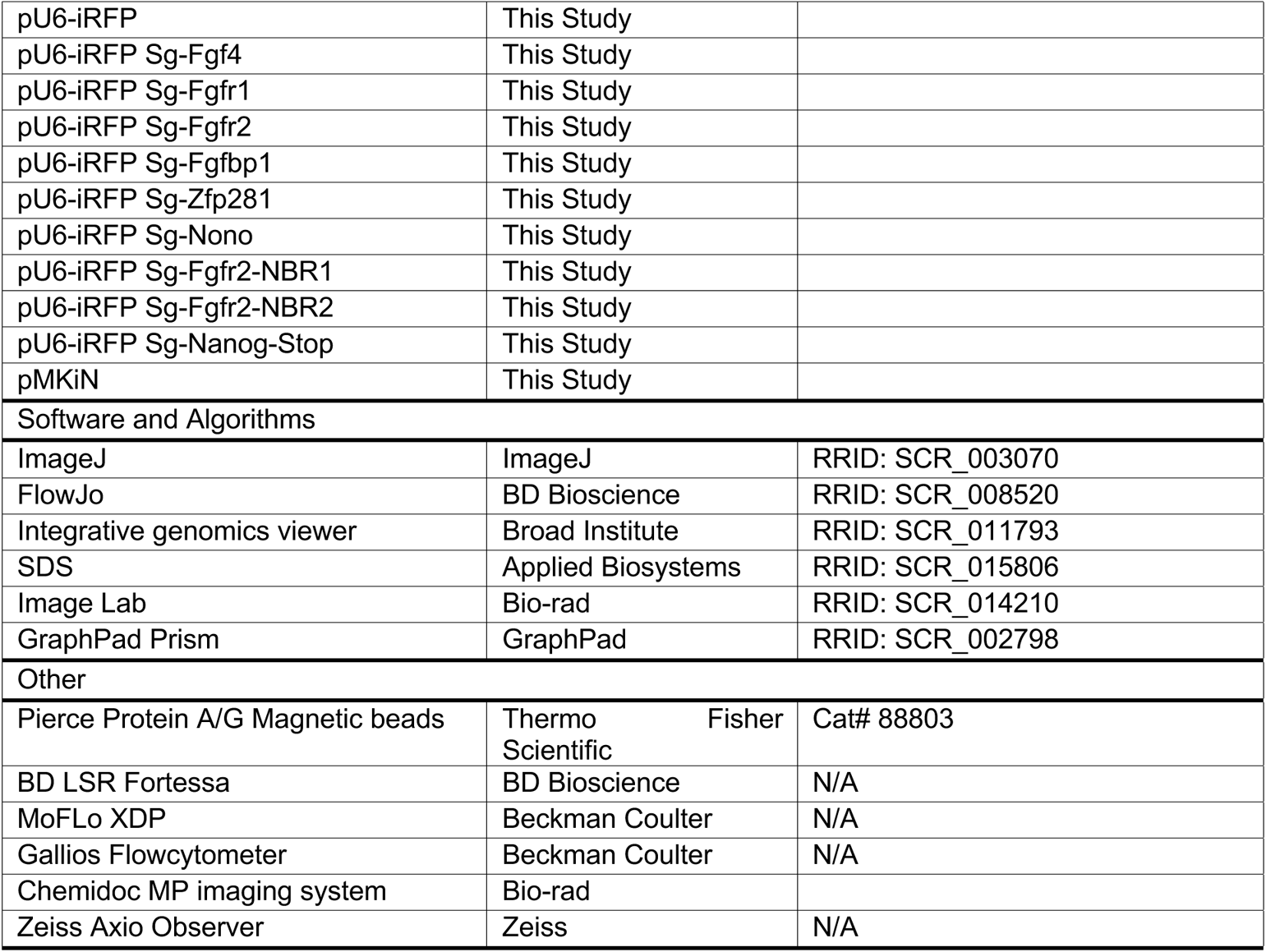

### Subhead. Methods

#### Cell Culture

The cell lines used in this study and their origin is depicted in Fig. S7. All the cells used in this study are derivatives of E14Tg2a ES. The cells were cultured as described earlier (5). 4-Hydroxytamoxifen (4-OHT), Doxycycline, and Cycloheximide were used at a concentration of 1 μg/ml, 1 μg/ml, and 100 μg/ml respectively. The TNERT and its derivative cell lines were treated with 4-OHT for 18 hrs except when indicated. TDiN and EDiN were treated with Doxycycline for 48 hrs unless indicated. CHIR99021 (CHIR, PD0325901 (PD), and SU5402 were used at 3 μM, 1 μM, and 2 μM, respectively, except when indicated. FGF4 and FGFBP1 were used at 50ng/ml concentration. The cells were cultured in Serum+ LIF (SL), SL+ PD (SLPD), SL+CHIR (SLCHIR), SL+SU5402 (SLSU5402), SL + PD +CHIR (SL2i) and N2B27+LIF+PD+CHIR (2iL) for at least 2 passages before treating with either 4-OHT or Doxycycline.

#### ELISA Assay

The condition media from the cell lines was collected at the respective time points. 100 µl of the media was coated per well of 96 wells of ELISA plate by incubating overnight at 4°C. The wells were washed thrice with PBS containing 0.05% Tween-20 and blocked with PBS containing 2% BSA for one hour at room temperature. The wells were washed once with PBS and incubated with the appropriate primary antibody (1:100) for one hour. Washed thrice with PBST, an appropriate HRP-labeled secondary antibody was hybridized for one hour at room temperature. The wells were washed thrice with PBST and incubated in substrate solution OPD (o-phenylenediamine dihydrochloride) 3mg/ml with 6 µl/ml H2O2) for 30 min in dark. The reaction was stopped by using 2N H2SO4. The absorbance was measured at 492 nm in Power wave XS2 (Bio Tek instruments).

---

**Fig. S1.**
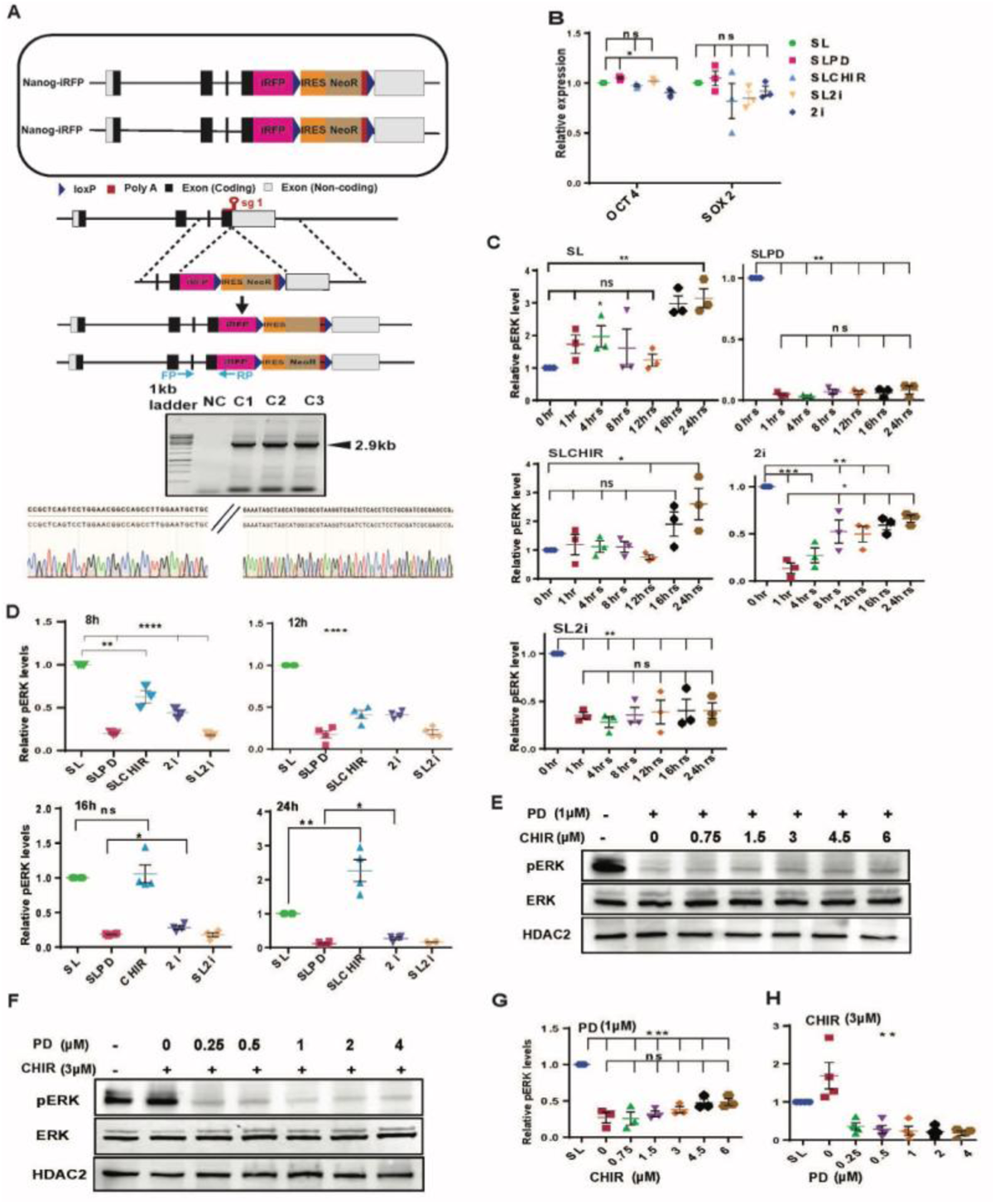
Residual MEK1/2 activity in the ground state prevents complete derepression of *Nanog*. (A) (top) Schematic depiction of NiRFP2A with both alleles of *Nanog* fused to iRFP coding sequences in frame with last coding sequence. (Middle) CRISPR mediated knock-in strategy of the iRFP-loxP-IRES-NeoR-loxP cassette into *Nanog* locus. The sgRNA includes the stop codon of the *Nanog* gene. The location of the genotyping primers (FP/RP) for the knock-in is marked by the arrows. (Bottom) Genotyping of the NiRFP2A clones, a 2.9 kb band is amplified only in the knock-in clones as one of the primers is complementary to a sequence outside the left homology arm and the other primer is complementary to the iRFP sequence. (B) Relative quantification of OCT4 and SOX2. The expression is normalized relative to HDAC2 levels and expression levels of OCT4 and SOX2 in SL (n>=3). (C, D) Relative pERK expression levels in indicated time points and treatments (n=3). (E) Western blot of pERK and ERK in 1 *μ*M PD and increasing concentrations of CHIR in SL media. (F) Western blot of pERK and ERK in 3 *μ*M CHIR and increasing concentrations of PD in SL media. (G, H) Relative pERK expression levels in indicated concentrations of CHIR and PD respectively (n>=3). All error bars in the figure represent s.e.m..

**Fig. S2.**
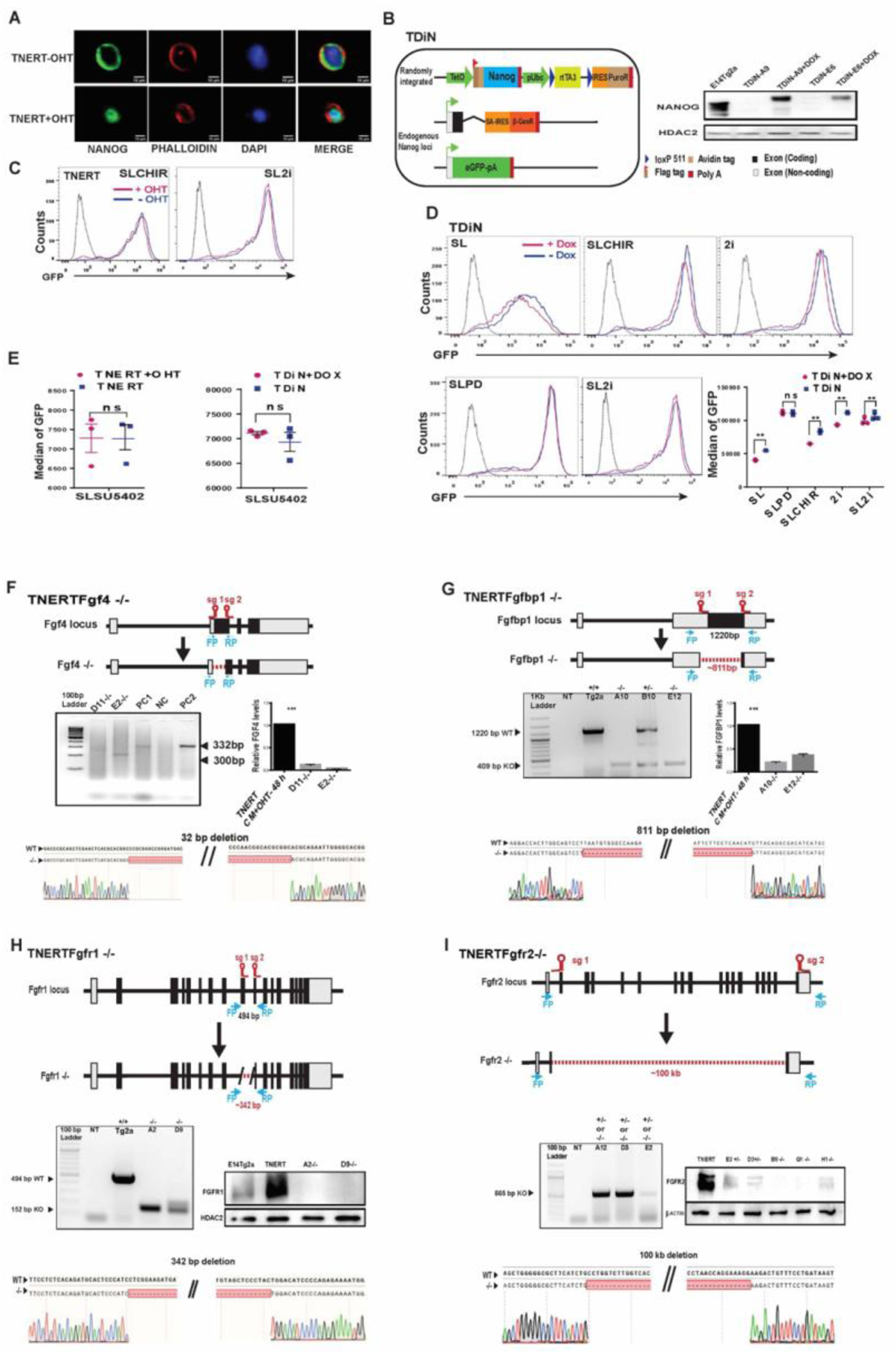
FGF autocrine signaling pathway components are essential for *Nanog* autoregulation. (A) Immunofluorescence of NANOG in TNERT cells after 30 mins treatment with or no OHT. Scale bars represent 10μM. (B) (left) Schematic of Doxycycline inducible TDiN cell line, generated by the introduction of a Tetracycline inducible Flag-Avi-NANOG (FaNANOG) transgene in Tβc44Cre6 cell line. (Right) Western blot of FaNANOG in TDiN cells after 48 hrs treatment with or no Doxycycline. (C) FACS profiles of TNERT treatment with or no OHT in SLCHIR and SL2i. D) FACS profiles of TDiN cell line in SL, SLCHIR, 2i, SL2i, and SLPD, (bottom right) *Nanog*:GFP population median of TDiN in indicated treatments (n=3). (E) *Nanog*:GFP population median of TNERT, NsGFPDiN, and TDiN treated with SU5402, with OHT/Doxycycline or no OHT/Doxycycline. (F) (top) CRISPR-based knock-out strategy using paired sgRNA to knock-out *Fgf4* in TNERT. The sgRNAs are positioned at the beginning and the near end of exon II. The deletion results in the loss of the start codon and a part of the coding region in exon II. The dotted line represents the deleted region of the gene. FP and RP represent the relative positions of the genotyping primers. (Middle left) genotyping PCR of TNERTFgf4−/− clones. The WT allele gives an amplicon of 332 bp and the knock-out allele a smaller amplicon by 32 bps or more. (Middle right) The relative abundance of FGF4 in media of TNERT and TNERTFgf4−/− clones 48 hrs after OHT treatment. (Bottom) Chromatogram of the TNERTFgf4−/− clones showing the sequences at the junction of the deletion. (G) (top) Schematic of the gene structure of *Fgfbp1* and the relative positions of the two sgRNAs used for paired sgRNA knock-out strategy. One sgRNA is complimentary to 5’UTR and the other to the 3’ end of the coding region of the only exon. (Middle left) Genotyping PCR showing a WT amplicon of 1220 bps and an amplicon around 400 bps in case of deletion. (Middle right) The relative abundance of FGFBP1 in media of TNERT and TNERTFgfbp1−/− clones 48 hrs after OHT treatment. (Bottom) Chromatogram of the TNERTFgfbp1−/− clone showing the sequences at the junction of the deletion. (H) Strategy for knock-out of *Fgfr1* in TNERT cells. The schematic depicts the gene structure of *Fgfr1* with the relative positions of the two sgRNAs. One sgRNA targets the 3’end of Intron 8 and the other exon10. (Middle left) Genotyping PCR shows a WT allele amplicon at 494 bp and a knock-out allele with smaller amplicons around 150 bp. (Middle right) Western blot analysis of FGFR1 in TNERT and TNERTFgfr1−/− clones. (Bottom) chromatogram showing the sequence of the deleted region. (I) A paired sgRNA strategy to knock-out *Fgfr2* in TNERT. (Top) The schematic represents the *Fgfr2* gene structure, with relative positions of the sgRNAs. One sgRNA target exon2 and the other sgRNA targets the coding region of the last exon approximately 100 kb apart. The dotted line represents the region of deletion in the *Fgfr2* gene. (Middle left) PCR genotyping shows a 665 bp amplicon when at least one allele of *Fgfr2* is deleted. This genotyping strategy cannot distinguish between +/− and −/− genotypes. (Middle right) Western blot analysis of FGFR2 protein in the *Fgfr2* targeted clones distinguishing the +/− and −/− clones. (Bottom) chromatogram represents the sequence of the genotyping amplicon indicating the exact sites of deletion. All error bars in the figure represent s.e.m.

**Fig. S3.**
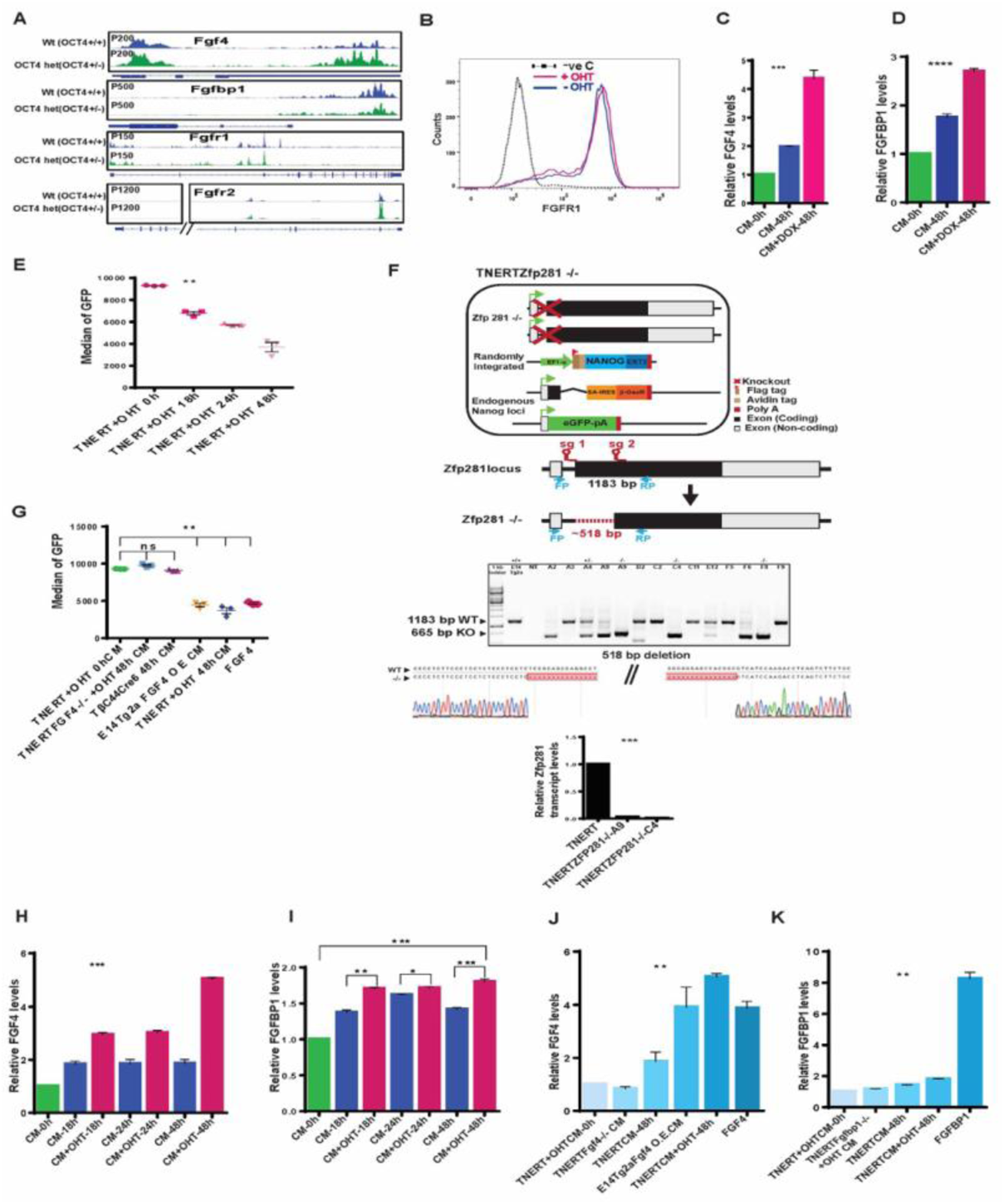
*Nanog* enhances expression of FGF autocrine signaling pathway components. (A) Browser tracks of NANOG enrichment in Fragment Per Kilobase of transcripts per Million (FPKM) in Oct4+/+ cells (normal NANOG levels) and Oct4+/− cells (higher NANOG levels) (Karwacki-Neisius et al., 2013) at *Fgf4, Fgfbp1, Fgfr1*, and *Fgfr2* loci. (B) Histogram of FGFR1 expression on the cell surface analyzed by immunostaining and FACS of fixed but unpermeabilized TNERT cells treated with (red) or no OHT (blue). (C, D) ELISA-based relative quantification of FGF4 and FGFBP1in media from EDiN cells cultured with or no Doxycycline (n=3). EDiN cell was generated by introducing a Doxycycline inducible Flag-Avi-NANOG transgene in E14Tg2a cells. (E) *Nanog*:GFP population median of Tβc44Cre6 treated with OHT induced conditioned media collected after different time points (n=3). (F) (top) Schematic of TNERTZfp281−/− cells, (upper-middle) CRISPR based paired guide knock-out strategy indicating the relative position of the sgRNAs, FP and RP indicate the genotyping primers. (Lower middle) Genotyping PCR indicating +/− and −/− clones. (Bottom) The sequencing chromatogram of the deleted region confirms the exact site of deletion, followed by RT-qPCR analysis of the *Zfp281* transcripts. (G) *Nanog*:GFP population median of Tβc44Cre6 treated with conditioned media from TNERT+OHT 0 hrs, TNERTFGF4−/− +OHT 48 hrs, Tβc44Cre6 48 hrs, E14Tg2a-FGF4-OE (overexpression) 48 hrs, TNERT+OHT 48 hrs and 50ng/ml FGF4 (n=3). (H, I) ELISA-based relative quantities of FGF4 and FGFBP1 in media from TNERT after 18, 24, and 48 hrs of OHT treatment (n=3). (J) ELISA-based relative quantities of FGF4 in conditioned media from cell lines - TNERT+ OHT 0 hrs, TNERTFGF4−/− + OHT 48 hrs, E14Tg2a-FGF4-OE 48 hrs (overexpression), TNERT−/+OHT 48 hrs, and 50ng/ml FGF4 (n=3). (K) ELISA-based relative quantities of FGFBP1 in conditioned media from various cell lines - TNERT+ OHT 0 hrs, TNERT-Fgfbp1−/− 48 hrs +OHT, TNERT 48 hrs −/+ OHT, and 50 ng/ml FGBP1 (n=3). All error bars in the figure represent s.e.m.

**Fig. S4.**
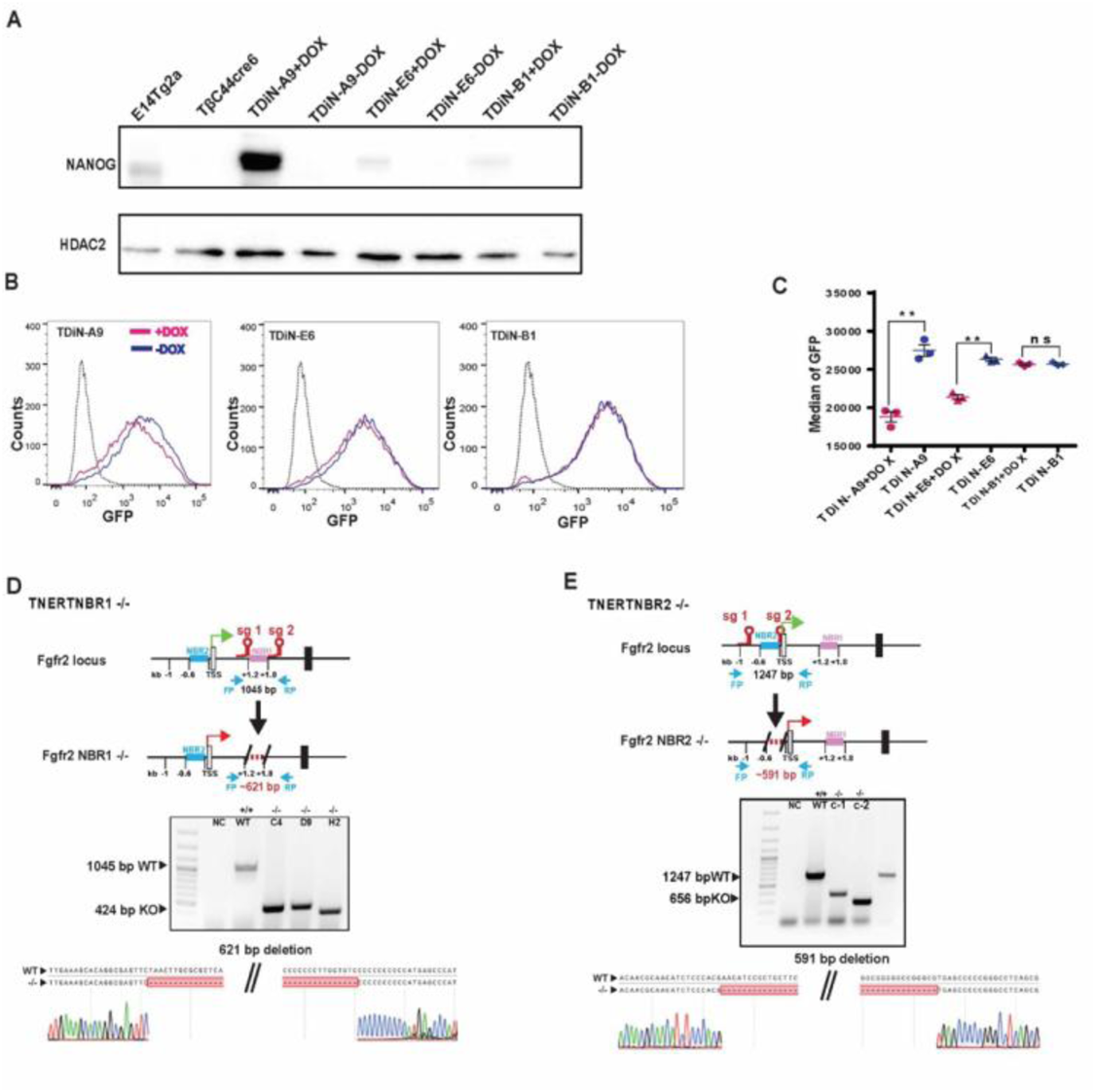
NANOG induced FGFR2 triggers autoregulation predominately in the ES cell population with higher *Nanog* expression. (A) Western blot analysis of Flag-Avi-NANOG in different clones of TDiN treated with or no Doxycycline showing different levels of NANOG expression relative to E14Tg2a. The clones show different levels of expression Flag-Avi-NANOG upon Doxycycline treatment. (B) FACS profiles of *Nanog*:GFP in TDiN clones treated with or no Doxycycline. (C) *Nanog*:GFP population median of TDiN clones (n=3). (D) (top) Schematic of strategy for deletion of NANOG Binding Region 1 (NBR1) in TNERT indicating the position of the NBR1 and the relative position of the sgRNA pair. The sgRNAs are complementary to sequences around 1.2 kb and 1.8 kb downstream of TSS. FP and RP indicate the relative position of the primers for genotyping. (middle) Genotyping of TNERT NBR1 knock-out clones. The WT shows an amplicon of 1045 bp, upon deletion around 600 bp sequence comprising multiple NANOG binding sites is deleted. (bottom) Sequence and chromatogram of the genotype PCR amplicon indicating the exact sequence of the junction of deletion in TNERTNBR1−/− clone. (E) Schematic of strategy for deletion of NANOG Binding Region 2 (NBR2) in TNERT indicating the position of NBR2 and relative position of the sgRNA pair. The sgRNAs are complementary to sequences around TSS and 0.6 kb upstream of TSS of *Fgfr2*. (middle) Genotyping of the TNERTNBR2 knock-out clones. The WT shows an amplicon of 1247 bp. The knock-out would lead to deletion of around 690 bps and a smaller amplicon of around 650 bps. (bottom) Sequence and chromatogram of the PCR amplicon from TNERT knock-out clones showing the exact site of deletion in TNERT NBR2−/− clone. All error bars in the figure represent s.e.m.

**Fig. S5.**
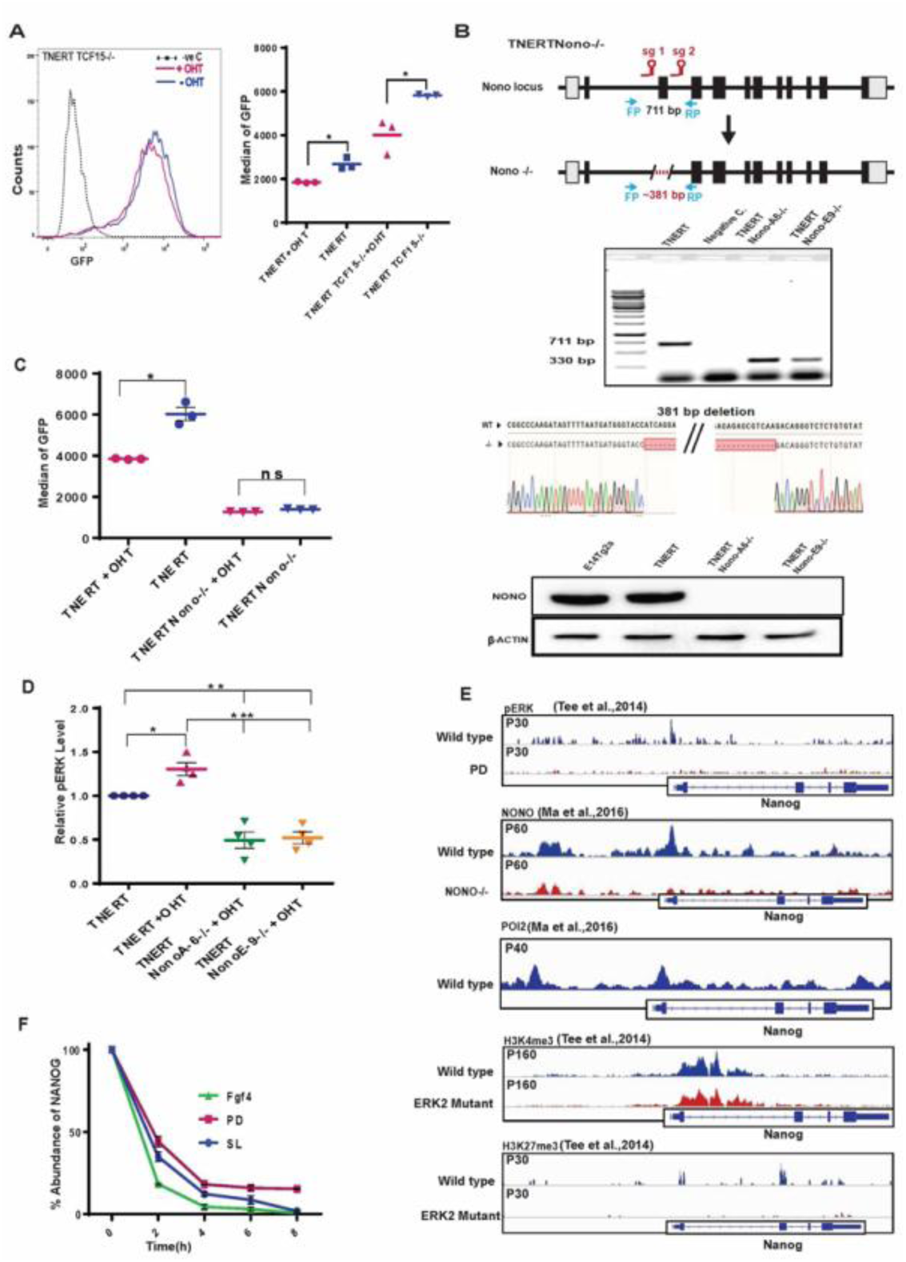
ERK interacts and recruits NONO to repress Nanog transcription. (A) (Left) FACS profile of TNERTTcf15−/− treated with or no OHT (n=3). (Right) *Nanog*:GFP population median of TNERT and TNERTTcf15−/−−/− treated with or no OHT (n=3). (B) A CRISPR-based knock-out strategy using paired sgRNA, to knock-out of *Nono* in TNERT cells. (Top) The schematic represents the mouse *Nono* gene structure with relative positions of the two sgRNAs flanking the second coding exon of *Nono*. FP and RP indicate the relative position of genotyping primers. The dotted line indicates the region of deletion in the *Nono* gene. (Middle) Genotyping PCR of the *Nono*−/− deletion in TNERT. The WT allele gave an amplicon of 711 bp and the deleted allele shows a smaller amplicon of 330 bp; followed by sequence and chromatogram indicating the deletion site (bottom) Western blot analysis of NONO protein in TNERT and TNERTNono−/− clones. (C) *Nanog*:GFP population median of TNERT and TNERTNono−/− treated with or no OHT (n=3). (D) The relative abundance of pERK in TNERT treated with or no OHT and TNERTNono−/− with OHT (n=4). (E) Browser tracks of pERK, NONO, POL2, H3K4me3, H3K27me3 enrichment in Fragment Per Kilobase of transcripts per Million (FPKM) on *Nanog* gene (Ma et al., 2016; Tee et al., 2014). (F) The relative abundance of NANOG after 0, 2, 4, 6, and 8 hrs of Cycloheximide (CHX) chase cultured in SL, PD, and FGF4 (n=3). All error bars in the figure represent s.e.m.

**Fig. S6.**
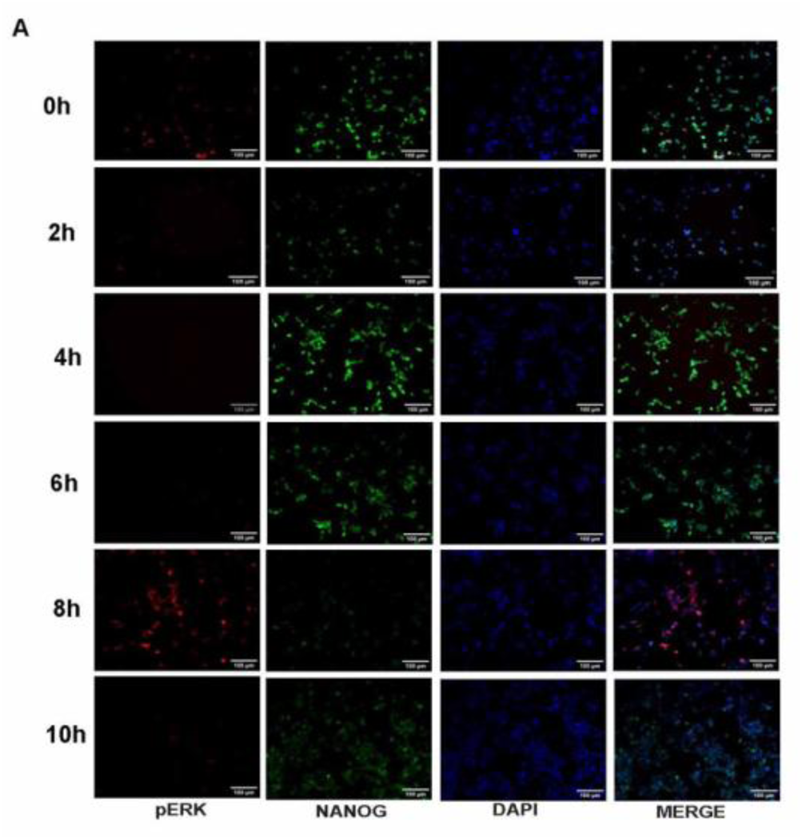
NANOG regulates ERK signaling dynamics and heterogeneity. (A) Immunocytochemistry at different time intervals of culture of NiRFP2A ESCs after sorting of 10% of *Nanog*-high cells. pERK (red), NANOG (green) and nuclear stain (blue). pERK and NANOG were detected in 10% *Nanog*-high cells immediately after sorting. Some of the cells expressed both pERK and NANOG. pERK decreased drastically within 2 hrs of culture. pERK1//2 was lowest at 4 hrs with concomitant high expression of NANOG. pERK expression was increased by 8 hrs coinciding with decreased NANOG. The pERK expression decreased by 10 hrs with increase in NANOG.

**Fig. S7:**
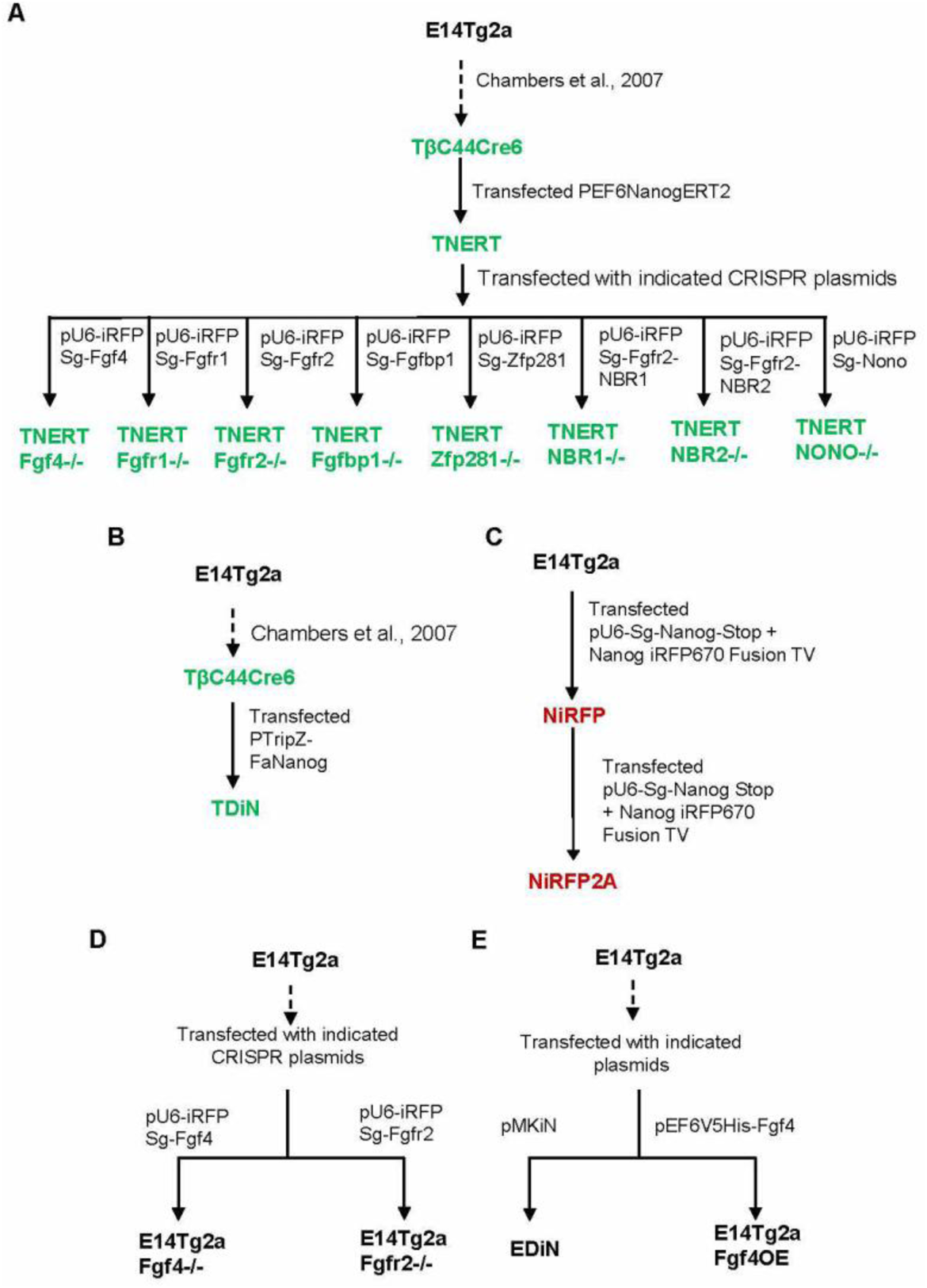
A pedigree chart of cell lines used in this study. (A) Flow chart illustrating the lineage and process of generation of TNERT and generation of knock out cells lines in TNERT background. (B) A flow chart describing derivation of TDiN. (C) A flow chart depicting derivation of NiRFP2A from E14Tg2a. (D) A flow chart depicting derivation of *Fgf4*−/− and *Fgfr2*−/− ES cell lines from E14Tg2a. (E) A flowchart depicting derivation of EDiN and *Fgf4*OE (over expression) ES cell lines from E14Tg2a.

**Table S1.**
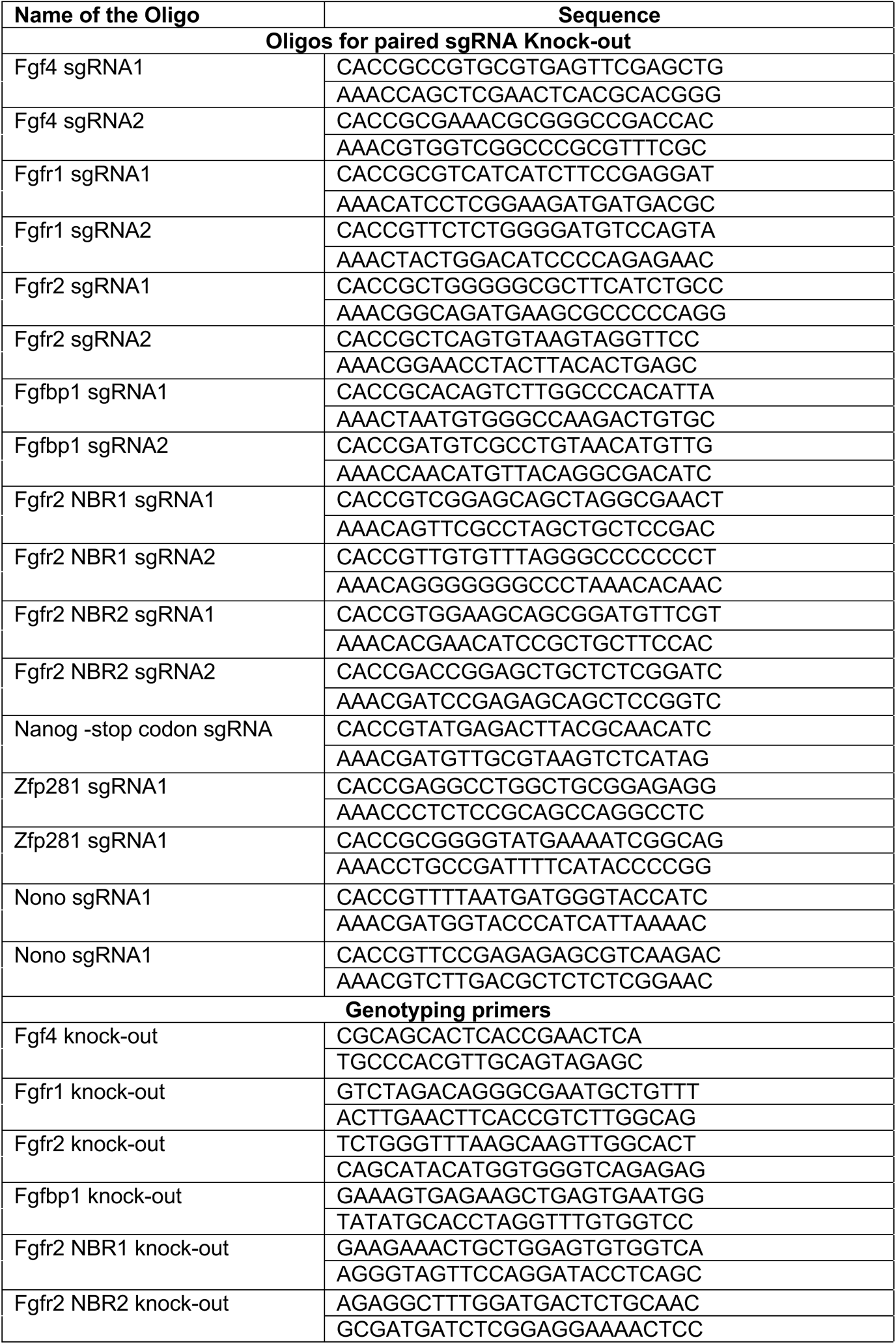
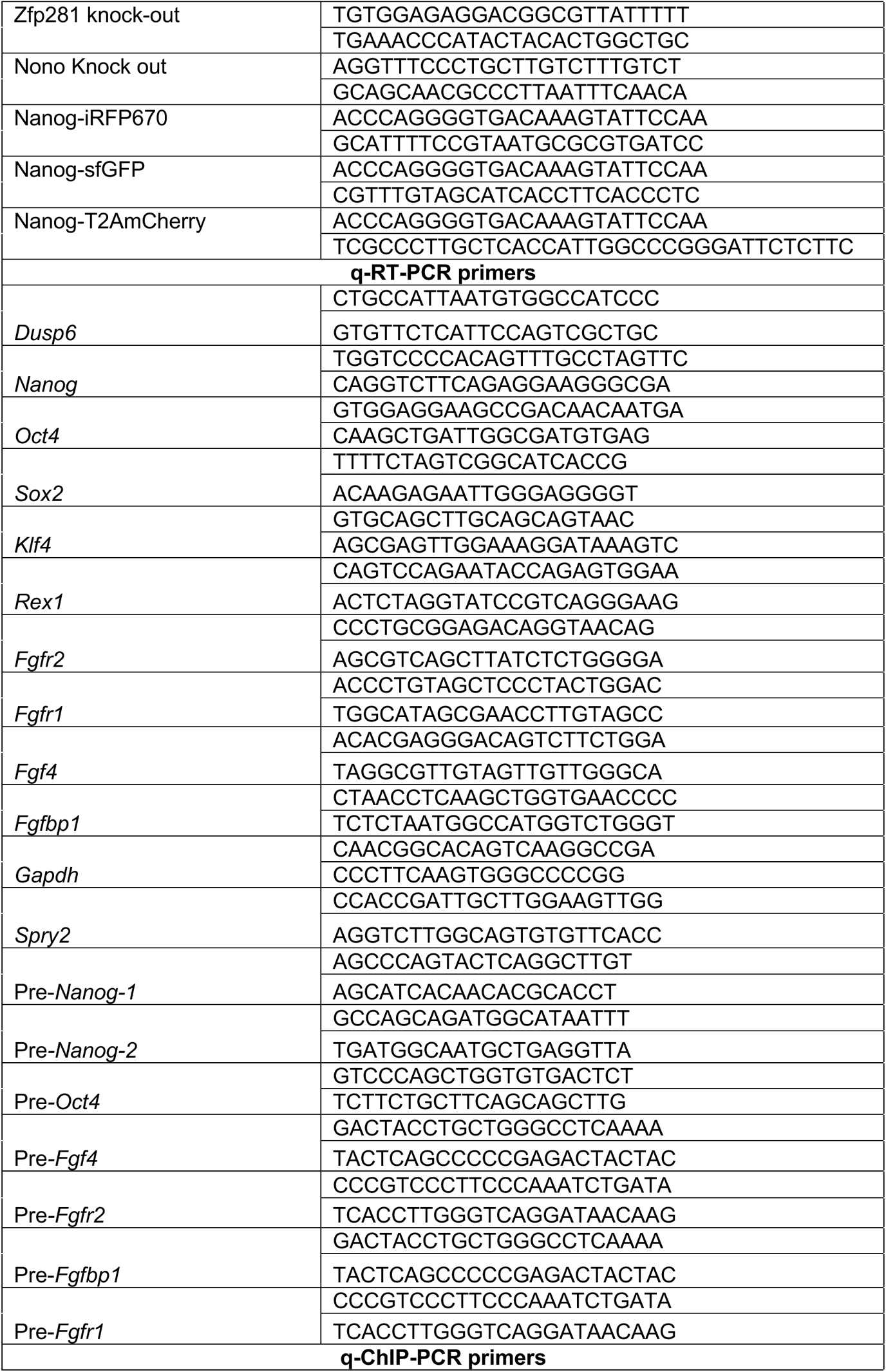
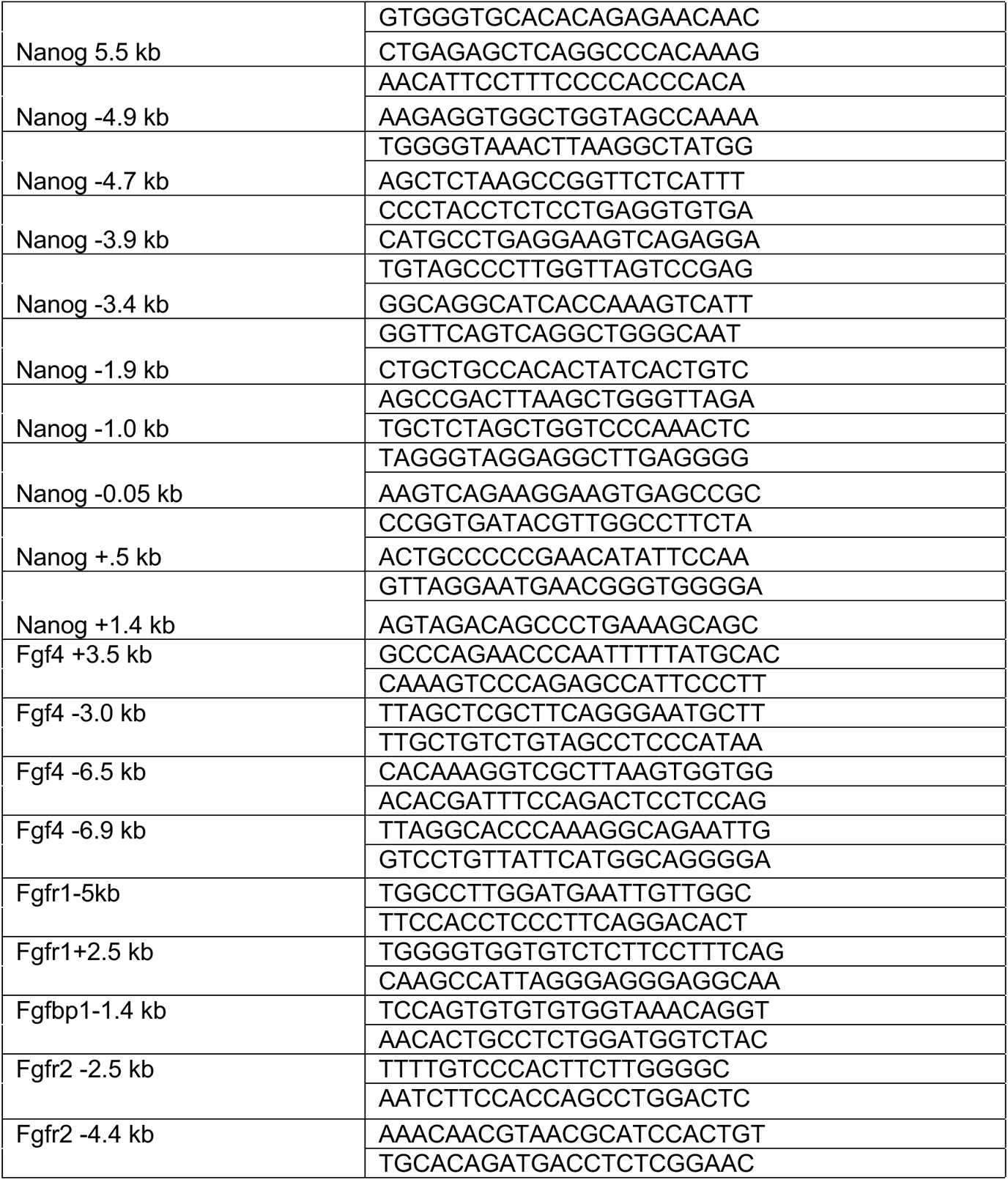
Oligonucleotide sequences used in this study.

## Notes

### Competing Interest Statement

The authors have declared no competing interest.

### Summary of Updates

To add the supplemental file and the resource tables

